# An Integrated Knowledge Graph and Network Medicine Pipeline for Drug Repurposing: Benchmarking Across Human Diseases and Application to Amyotrophic Lateral Sclerosis

**DOI:** 10.64898/2026.07.03.736387

**Authors:** Anqi Jiang, Jiajing Hu, Yusuf Abdulle, Oliver Pain, Alfredo Iacoangeli

## Abstract

Drug repurposing offers a practical strategy to identify new therapeutic uses for approved drugs, potentially reducing the time and cost associated with conventional drug development. We present a novel three-stage drug repurposing pipeline that integrates knowledge graph-based gene prediction, network-based drug-disease association analysis, and systematic classification of candidate drugs by therapeutic class. The pipeline integrates DGLinker to predict novel disease-associated genes, SAveRUNNER to identify drug repurposing candidates, and ATC Category Enrichment Analysis (ATCEA) to prioritise candidates by pharmacological class.

We benchmarked the pipeline across twelve diseases using DrugBank and MEDI2-HPS as validation resources. Utilising DGLinker-expanded disease-gene sets as input increased the number of predicted repurposed drugs, while overall discriminative performance remained stable across diseases (AUROC 0.71-0.77). Application of ATCEA consistently improved precision, F1-score, and specificity, while reducing recall, reflecting a conservative prioritisation strategy that contracts the candidate space while retaining pharmacologically coherent drug-disease candidates.

We further applied the pipeline to amyotrophic lateral sclerosis (ALS), a neurodegenerative disease with limited therapeutic options and performed a deeper literature-based validation of the results. Incorporation of DGLinker-predicted genes substantially increased the number of significant candidate drugs and uncovered enriched ATC categories not identified using known ALS genes alone, including antidepressants and antipsychotics. Moreover, several drugs with supporting evidence available in literature were identified only when DGLinker-predicted genes were used. Overall, 77 candidate drugs were prioritised within significantly enriched ATC categories, several of which are supported by previously published studies. To provide exploratory real-world support for these findings, we further evaluated candidate drugs in a longitudinal electronic health record (EHR) dataset of 2361 patients with ALS from King’s College Hospital. Although the number of evaluable drugs was limited due to sample size, the EHR analysis provided additional clinically relevant context for selected prioritised drugs and pharmacological classes.

Our pipeline demonstrates potential to accelerate drug repurposing by integrating complementary computational approaches to each step of the process, providing an end-to-end framework that showed robust performance across benchmarking experiments and use cases.

**Graphical abstract:** 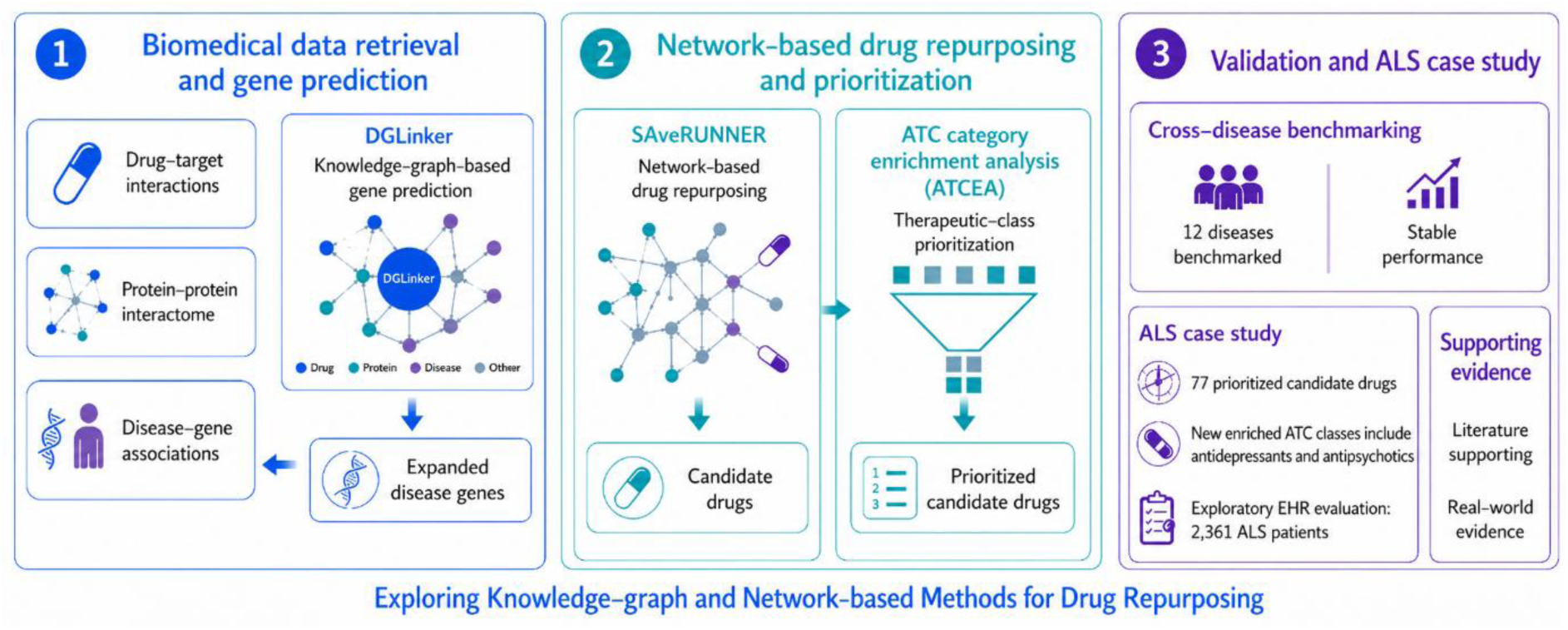

## Introduction

Drug repurposing is a promising strategy for identifying new therapeutic applications for existing approved compounds. By bypassing early-stage development and leveraging existing safety and pharmacokinetic data, this approach reduces both the time and cost of bringing treatments to market (Novac, 2013). It is especially relevant for rare diseases with limited therapeutic options, for which rapid treatment is greatly needed (Shah et al., 2021). Recent computational approaches to drug repurposing include machine learning, matrix factorization, deep learning, and network-based methods (Cai et al., 2023). Traditional machine learning models are efficient but often rely on manually engineered features and on assumed negative samples (i.e., drug-disease pairs with no known therapeutic association are treated as negatives). These requirements can introduce bias and hinder scalability (Nascimento et al., 2016; You et al., 2018; B. Yu et al., 2020). Matrix factorization methods address missing data but suffer from limitations such as narrow prediction range and convergence to local minima (Kim et al., 2023; Sadeghi et al., 2022; M. Wang et al., 2018). Deep learning frameworks excel at processing complex, large-scale data with automated feature learning, yet their interpretability, computational demands, and reliance on extensive labeled datasets constrain their applicability in understudied conditions (Cai et al., 2021; Meng et al., 2022; Sun et al., 2024; Zeng et al., 2019).

In contrast, network-based methods offer improved interpretability and flexibility. For example, SAveRUNNER (Fiscon et al., 2021) identifies candidate drugs based on proximity between drug targets and disease genes in the interactome, without requiring prior drug-disease associations. This independence improves its applicability to diseases lacking approved therapies or extensive annotation. Its utility has been demonstrated across diverse conditions, including Alzheimer’s disease, COVID-19, and several cancers, with predictions supported by pathway enrichment and literature-based validation (Brunetti et al., 2023; Fiscon et al., 2022; Sibilio et al., 2021), demonstrating practical value in drug repurposing. However, the performance of such methods greatly depends on our understanding of the disease genetics.

While currently annotated disease-associated genes have contributed significantly to understanding pathophysiology and to therapeutic targeting (Reay & Cairns, 2021), they often provide an incomplete picture due to our limited understanding of the genetic basis of most heritable human diseases. This limitation is particularly evident for complex neurodegenerative diseases, and it is well exemplified by Amyotrophic Lateral Sclerosis (ALS). Although known disease-causing mutations explain most familial ALS cases, and the heritability of the sporadic cases (approx. 90-95% of all cases) is estimated to be ∼60% (Al-Chalabi et al., 2010), known ALS genes can causally explain only ∼20% of all people with ALS (Cook & Petrucelli, 2019; Iacoangeli et al., 2025; Mehta et al., 2022; Van Daele et al., 2023). Given that drug repurposing methods rely on a set of disease-gene associations, and the observed correlation between strong genetic evidence and clinical success of drug targets (Nelson et al., 2015), expanding the gene-disease space using computational prediction methods may offer a valuable strategy for improving drug repurposing pipelines.

In recent years, several studies have incorporated expanded disease gene sets into drug repurposing workflows through gene prioritisation algorithms and machine learning models (Xu et al., 2022; Yan et al., 2021; Zhu et al., 2020). While these approaches demonstrate the feasibility of this strategy in drug discovery, most do not explicitly evaluate whether such genes offer added value compared to the use of known disease genes. As a result, the impact of novel predicted disease genes on drug prioritisation remains unclear.

Building on this foundation, we developed a new end-to-end drug repurposing pipeline that integrates both knowledge-based gene-disease and network-based drug-disease association prediction methods with the aim of exploring the impact of such a strategy for drug repurposing. The pipeline includes three main steps: 1) First it employs DGLinker (Hu et al., 2021), a knowledge graph-based tool selected for its high performance, flexibility, and ability to integrate diverse biological and phenotypic data sources (Bean et al., 2020; Hu et al., 2021). DGLinker generates an expanded set of genes associated with the target disease, incorporating both known disease genes from reference databases and novel computational predictions; 2) Then, it uses known disease genes, and an expanded gene set that includes novel disease-gene predictions from DGLinker, as input for SAveRUNNER, a network-based method that predicts repurposable drugs using proximity between drug targets and disease genes in protein interaction networks; 3) Finally, to further prioritise candidate drugs, we incorporated Anatomical Therapeutic Chemical (ATC) Category Enrichment Analysis (ATCEA), which aggregates predicted candidates by pharmacological class. Enriched ATC categories highlight drug classes that are overrepresented among network predictions, enabling pharmacologically consistent prioritisation of candidate drugs. Together, DGLinker, SAveRUNNER, and ATCEA form a modular and interpretable pipeline designed to improve the identification of repurposed drugs, particularly in biologically complex and therapeutically underserved diseases. To our knowledge, our pipeline addresses a methodological gap by providing a drug repurposing framework that integrates upstream computational disease gene predictions with network-based drug prioritisation and pharmacological class-level prioritisation.

In this article, we first describe the pipeline and present a validation experiment benchmarking its predictions using known disease-drug associations from DrugBank and MEDI2-HPS databases (Knox et al., 2024; Wei et al., 2013). We then apply the pipeline to a case study of ALS, a fatal and devastating neurodegenerative disease with a treatment landscape remarkably constrained. Indeed, Riluzole (Rilutek) remains the only widely available and routinely prescribed pharmacological treatment for ALS within the United Kingdom (Gallagher et al., 2021), and its clinical utility is limited by modest efficacy, extending survival by 2-3 months and primarily showing benefit in late-stage disease manifestation (Hinchcliffe et al., 2017). While a gene-targeted therapy, Tofersen, has received regulatory approval for a small genetically defined subset of ALS patients, effective therapeutic options for the broader ALS population remain extremely limited (Hamad et al., 2025). To further support the clinical relevance of the prioritised ALS candidate drugs, we also performed an electronic health record (EHR) evaluation using patient data.

## Methods

We developed a multi-stage drug repurposing pipeline combining disease-gene prediction, network-based drug-disease association analysis, and pharmacological enrichment. The pipeline integrates DGLinker for gene set expansion, SAveRUNNER for drug prediction based on network proximity, and ATC Category Enrichment Analysis (ATCEA) for drug prioritisation. The following sections detail the data sources, computational methods, and validation framework used (Figure 1).

**Figure 1.**
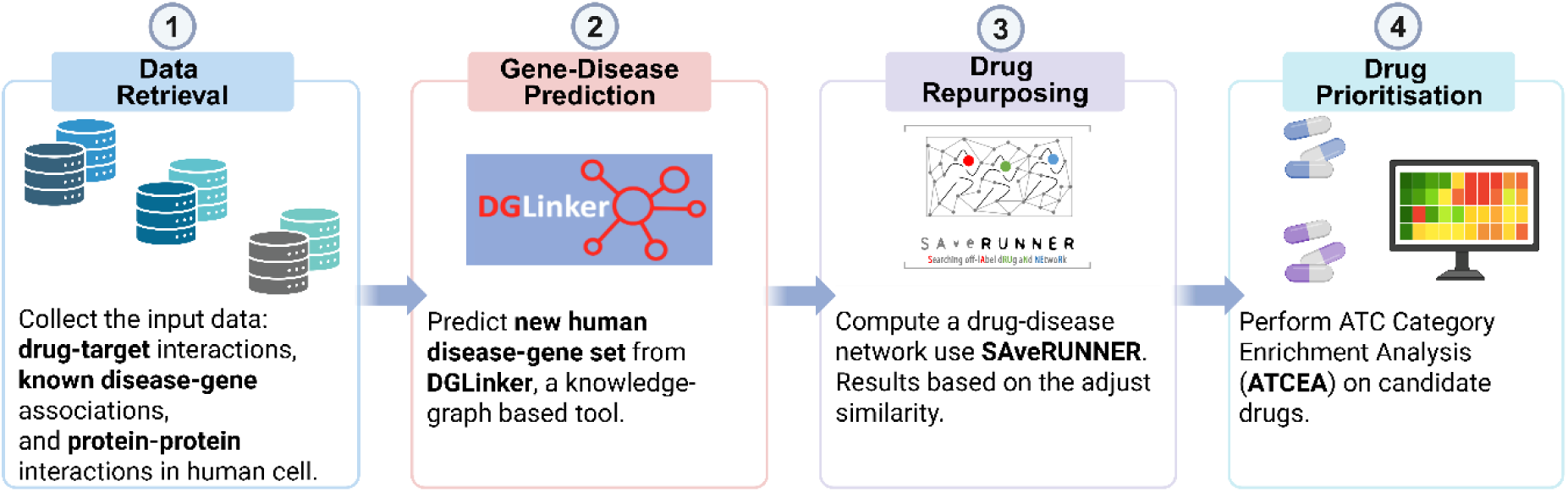
Overview of the drug repurposing pipeline workflow. *Created with* BioRender.com.

### Data Retrieval

We integrated multiple publicly available biomedical databases to construct the foundational networks used in the drug repurposing pipeline. These include drug-target, disease-gene, and protein-protein interaction datasets.

#### Drug-target interactions

We leveraged information from DrugBank v5.1.12 (Knox et al., 2024). Protein targets were mapped to their source genes using UniProt and standardized to NCBI Entrez Gene IDs (Ostell, 2012). Our database comprises 6,855 distinct drugs, forming 19,547 validated drug-target interaction pairs.

#### Disease-gene associations

Known disease-genes are sourced from the Comparative Toxicogenomics Database (CTD) [accessed on 30th Oct. 2024] (Davis et al., 2025), focusing on chemical-gene-disease relationships, and Phenopedia (W. Yu et al., 2010), documenting human gene-phenotype associations from genetic association studies. The combined database contains 6,964 distinct diseases, establishing 516,266 validated disease-gene associations. We also included a newly predicted disease-gene set from DGLinker (Hu et al., 2021), a knowledge-graph based tool for the prediction of novel human disease-gene associations. Application details are provided in the later “Gene-Disease Prediction” section.

#### Protein-protein interactions

data used in this analysis was obtained from the comprehensive human protein-protein interactome assembled by Cheng et al. (2018). To construct this interactome, the researchers integrated protein-protein interaction data from 15 databases, encompassing a wide range of experimental sources. The resulting interactome comprised 217,160 protein-protein interactions connecting 15,970 unique proteins.

### Gene-Disease Prediction

DGLinker (Hu et al., 2021) is a public webserver that integrates phenotypic information and biological data into a biomedical knowledge graph and applies machine learning to predict novel disease-associated genes. In our analysis, we selected the "Select phenotype(s)" option to input the phenotype term(s) of interest, allowing DGLinker to automatically retrieve genes associated with the phenotype from integrated databases, including DisGeNET, OMIM, ClinVar, and HPO. To construct the knowledge graph, we explicitly selected three types of databases provided by DGLinker: Gene-Disease Association (DisGeNet v7.0, Piñero et al., 2020), Gene Function (Gene Ontology v2021-02, Consortium et al., 2023), and Protein-Protein Interaction (IntAct v2021-04, del Toro et al., 2022). DGLinker subsequently trained a supervised classification model using this knowledge graph and ranked all human protein-coding genes based on their similarity scores relative to the input phenotype genes. The results included a downloadable text file detailing each gene’s rank, similarity score, and association type ("Input" or "Predicted"). For subsequent analyses, we exclusively selected genes labeled as "Predicted," thereby generating a new candidate gene set.

### Drug Repurposing

SAveRUNNER is an R-based tool for drug repurposing that applies a network-based inference strategy based on network medicine principles (Fiscon & Paci, 2021). It models diseases as localized perturbations within the human protein-protein interaction network and identifies candidate drugs by assessing the topological proximity between drug targets and disease-associated genes. The algorithm computes a network proximity score, evaluates its statistical significance against a random background, and further prioritises associations using a modularity-adjusted similarity measure. The final output is a weighted bipartite network of drug-disease links, ordered by their adjusted similarity scores.

#### Input

SAveRUNNER requires three primary data sets that we obtain from Data Retrieval step: drug-target, disease-gene, and gene-gene interactions.

#### Algorithm

The algorithm comprises four main stages: (1) computation of network-based drug-disease proximity and its statistical significance, (2) selection of statistically significant drug-disease associations, (3) computation of similarity measures and cluster detection, and (4) adjustment and normalization of network similarity. Steps 1-2 yield a proximity-based bipartite network in which nodes represent drugs and diseases, and edges denote statistically significant proximity between drug targets and disease-associated genes. Steps 3-4 then produce a similarity-based bipartite network that captures normalized similarity relationships within the same network neighborhoods. Although SAveRUNNER is fundamentally a proximity-based network method, in this study we use the statistical significance (p-value) of drug-disease proximity as the primary statistic for downstream validation and candidate selection, because p-values provide a normalized and comparable measure across diseases and databases.

#### Output

SAveRUNNER constructs a weighted bipartite network that elucidates the intricate connections between drugs and diseases. The edges in this network were established when the proximity between drug targets and disease-associated genes within the interactome network was statistically significant after Bonferroni correction (Bonferroni-adjusted p-value ≤ 0.05). The weight assigned to each edge represents the strength of the drug-disease association, derived from an adjusted similarity measure computed by the algorithm.

### Drug Prioritisation

The ATC Classification System is a hierarchical system that categorizes drugs according to their therapeutic, pharmacological, and chemical properties. ATC Category Enrichment Analysis (ATCEA) identifies overrepresented ATC categories within a set of drugs. In this study, ATCEA was used to prioritise SAveRUNNER-predicted drugs by detecting enriched pharmacological classes at the third ATC level. Drugs that had an association with a target disease in the drug-disease network from SAveRUNNER were mapped to third-level ATC categories, and ATCEA was assessed using the Wilcoxon-Mann-Whitney test, with multiple testing controlled using the Benjamini-Hochberg false discovery rate procedure (BH procedure). Drugs were prioritised if they belonged to significantly enriched ATC categories (BH-adjusted p-value ≤ 0.05) and showed Bonferroni-significant SAveRUNNER drug-disease associations. Further methodological details are provided in Supplement Document 1.

### Pipeline Validation

Our evaluation focuses on twelve diseases: ten diseases from the MEDI2 dataset (Wei et al., 2013) with the highest-precision drug-disease associations, together with Alzheimer’s disease and depressive disorder, given their relevance to ALS, the case-study disease described later in the Results. Alzheimer’s disease is a neurodegenerative disease that exhibits genetic and phenotypic correlations with ALS (Spargo et al., 2024; X. Wang et al., 2014). Additionally, since depressive disorders are present in 4-56% of ALS patients (Atassi et al., 2010), understanding drug associations in this context was valuable. Schizophrenia, another neuropsychiatric disorder that presents shared genetics with ALS was already considered as part of the top 10 diseases in MEDI2. We evaluated the predictive performance of our pipeline by comparing its output against reference drug-disease associations from two established databases: DrugBank v5.1.12 (Knox et al., 2024) and the high-precision subset of MEDI2 (MEDI2-HPS, released in 2020) (Wei et al., 2013). For each disease, the full evaluation universe was defined as the input drugs set, comprising all drugs subjected to repurposing analysis. All validation metrics were computed with respect to this evaluation universe. True positives (TP) were defined as drugs in this universe with approved indications for the target disease in DrugBank or MEDI2-HPS, while all remaining drugs were treated as negatives. In order to evaluate the performance of our pipeline, we estimated the Area Under the Receiver Operating Characteristic Curve (AUROC) using the pipeline-derived p-values as scores to rank drugs for a given disease, with known associations in the reference databases treated as positives and all others treated as negatives. We also computed a broad set of metrics, including precision, recall, F1-score, accuracy, and specificity, before and after using ATCEA as a post hoc filter on drug-disease associations. Furthermore, we quantified candidate drugs retention as the ratio of retained to initial candidate drugs for each disease. We did not use AUROC to evaluate the predictive performance of ATCEA filter because the filter operates by restricting the predictions to a subset of drugs, meaning that negatives outside the enriched classes are automatically excluded; this changes the score composition of the evaluation set and makes AUROC values difficult to interpret or directly compare with those from the unfiltered predictions.

To evaluate the added value of each pipeline component, we compared its predictive performance across three validation scenarios, which differed in their drug-target interaction network and disease-gene association inputs, before and after applying the ATCEA filter: 1) *Old network with known disease genes*, replicating the original SAveRUNNER study using the same drug-target network and only known disease-gene associations from reference databases; 2) *Updated network with known disease genes*, using the updated drug-target network and known disease-gene associations built from the latest biological databases (as described above); and 3) *Updated network with DGLinker-expanded gene set*, based on the 2) network but incorporates an expanded disease-gene set that includes both known and DGLinker-predicted genes. This design allows us to separately assess the effects of using more recent interaction data and expanding the disease-gene set on drug repurposing performance.

### EHR Evaluation of ALS Candidate Drugs

To provide exploratory real-world support for the ALS candidate drugs prioritised by our pipeline, we conducted an evaluation using a longitudinal EHR dataset comprising 2361 patients with ALS from King’s College Hospital between 2003 and 2023. The dataset integrated free-text clinical notes, laboratory observations, medication and diagnostic orders, and outpatient attendance records, linked at patient level using pseudonymised unique identifiers. Confirmed ALS cases were identified using specialist clinic documentation, diagnosis-related text patterns in clinical notes, and concept-level natural language processing methods implemented with MedCAT (Kraljevic et al., 2021).

For the ALS analysis, the diagnosis event was defined as the first *Neurology Motor Nerve Clinic Letter*. Candidate drugs identified in the ALS case study were mapped to prescription records, and only drugs observed in the EHR were considered for downstream exposure and survival analyses. We assessed two complementary survival settings: a pre-diagnosis matched survival analysis and a post-diagnosis landmark survival analysis. In the pre-diagnosis analysis, exposed patients were matched to patients without recorded pre-diagnosis exposure to the same drug, using age at diagnosis, sex, concurrent medication count, and calendar year of diagnosis. The matching setting used a 1:5 ratio with an age caliper of ±5 years. To ensure statistical robustness, only drugs with more than four exposed patients were included in the final analysis. In the post-diagnosis analysis, only patients who survived at least 0.3 years after diagnosis were included, and survival was measured from the 0.3-year landmark onward. This landmark design was used to reduce immortal time bias, which can arise because patients must survive long enough after diagnosis to receive post-diagnosis prescriptions (Gleiss et al., 2018). Survival associations were summarized using Hazard Ratios (HRs) and corresponding confidence intervals, Kaplan-Meier curves were plotted for visual comparison of survival between exposed patients and matched controls, and differences in survival distributions were assessed using the log-rank test. To account for multiple testing across candidate drugs, log-rank p-values were adjusted using the BH procedure. In addition, EHR-observable drugs were grouped according to third-level ATC categories, and class-level survival analyses were conducted to examine whether ATC categories enriched in the ALS case study showed corresponding survival patterns in the EHR data.

## Results

In this study, we developed a drug repurposing pipeline and benchmarked its performance across multiple diseases using approved drug-disease associations as reference standards. We then applied the pipeline to ALS, a fatal neurodegenerative disease with limited therapeutic options, to investigate the potential relevance of the predicted candidate drugs.

### Pipeline Validation

#### AUROC Performance and Candidate Space Variation

The drug repurposing pipeline performance in three validation scenarios was assessed using AUROC based on DrugBank and MEDI2-HPS validation databases (Figure 2). Additionally, the number of significant candidate drugs and recovered known drug-disease associations were examined to characterize changes in candidate space (Table 1).

**Figure 2.**
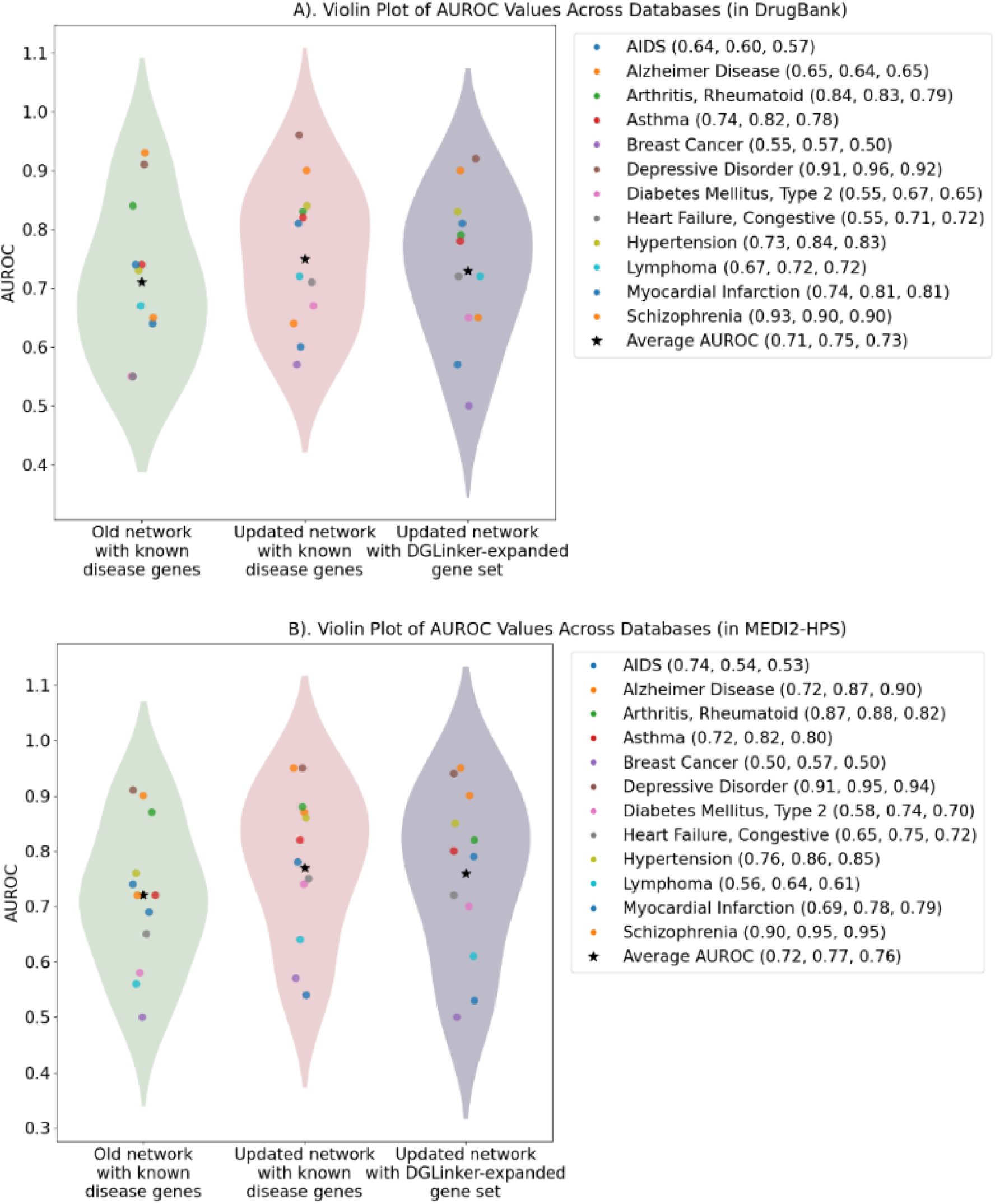
AUROC comparison of drug repurposing results in A) DrugBank and B) MEDI2-HPS. AUROC values for the three scenarios are reported between brackets in the figure legends.

**Table 1.** Candidate and matching drugs identified across validation scenarios databases. “Number of known drugs” denotes reference drugs with approved indications for the target disease in DrugBank or MEDI2-HPS that are present in the corresponding evaluation universe for each validation scenario. As the input drug set varies across scenarios, this number may differ accordingly.

| Disease | Old network with known disease genes |  |  |  |  |  | Updated network with known disease genes |  |  |  |  |  | Updated network with DGLinker-expanded gene set |  |  |  |  |  |
| --- | --- | --- | --- | --- | --- | --- | --- | --- | --- | --- | --- | --- | --- | --- | --- | --- | --- | --- |
|  | Candidate drugs | Input genes | Number of known drugs |  | Matching drugs (true positives) |  | Candidate Drugs | Input genes | Number of known drugs |  | Matching drugs (true positives) |  | Candidate drugs | Input genes | Number of known drugs |  | Matching drugs (true positives) |  |
|  |  |  | Drug Bank | MEDI2-HPS | Drug Bank | MEDI2-HPS |  |  | Drug Bank | MEDI2-HPS | Drug Bank | MEDI2-HPS |  |  | Drug Bank | MEDI2-HPS | Drug Bank | MEDI2-HPS |
| AIDS | 21 | 1055 | 29 | 22 | 1 | 1 | 21 | 1056 | 54 | 30 | 2 | 1 | 74 | 1886 | 54 | 30 | 0 | 0 |
| Alzheimer's disease | 125 | 1749 | 16 | 9 | 4 | 2 | 177 | 1787 | 46 | 9 | 4 | 3 | 170 | 2227 | 46 | 9 | 2 | 2 |
| Arthritis, Rheumatoid | 58 | 1028 | 53 | 49 | 4 | 5 | 100 | 1102 | 84 | 49 | 13 | 8 | 144 | 2190 | 84 | 49 | 3 | 4 |
| Asthma | 103 | 1315 | 58 | 60 | 4 | 5 | 144 | 1328 | 84 | 60 | 6 | 3 | 275 | 1906 | 84 | 60 | 8 | 8 |
| Breast cancer | 100 | 1788 | 51 | 29 | 2 | 0 | 73 | 2809 | 80 | 29 | 3 | 0 | 105 | 3494 | 80 | 29 | 0 | 0 |
| Depressive disorder | 268 | 626 | 22 | 49 | 17 | 41 | 385 | 649 | 25 | 49 | 18 | 38 | 465 | 2019 | 25 | 49 | 16 | 34 |
| Diabetes mellitus, Type 2 | 139 | 3654 | 32 | 70 | 0 | 2 | 151 | 3747 | 38 | 72 | 0 | 1 | 163 | 4651 | 38 | 72 | 0 | 1 |
| Heart failure, Congestive | 161 | 265 | 65 | 69 | 9 | 17 | 238 | 265 | 81 | 70 | 12 | 12 | 460 | 1205 | 81 | 70 | 7 | 8 |
| Hypertension | 159 | 2121 | 121 | 110 | 17 | 19 | 177 | 2176 | 152 | 111 | 22 | 17 | 184 | 2176 | 152 | 111 | 22 | 18 |
| Lymphoma | 77 | 1174 | 50 | 65 | 3 | 2 | 107 | 1178 | 86 | 65 | 2 | 3 | 78 | 2410 | 86 | 65 | 1 | 0 |
| Myocardial Infarction | 146 | 930 | 50 | 95 | 7 | 11 | 221 | 975 | 60 | 97 | 12 | 11 | 240 | 1684 | 60 | 97 | 7 | 10 |
| Schizophrenia | 299 | 1809 | 36 | 30 | 31 | 24 | 365 | 1875 | 62 | 30 | 34 | 24 | 369 | 1875 | 62 | 30 | 36 | 24 |

The average AUROC values were similar across the scenarios (0.71-0.75 in DrugBank and 0.72-0.77 in MEDI2-HPS). Depressive disorder and schizophrenia consistently showed high discriminative performance (AUROC > 0.90), whereas breast cancer showed the lowest AUROC values across settings (AUROC < 0.57). Updating the network and incorporating DGLinker-expanded gene sets resulted in only modest, disease-specific AUROC changes, with no systematic improvement and an overall stable AUROC distribution. For example, AUROC increased for Alzheimer’s disease in MEDI2-HPS, whereas slight decreases were observed for AIDS in both validation datasets.

Beyond AUROC performance, we further examined how network updates and gene expansion influenced the number of significant candidate drugs and the identification of matching drugs (true positives) (Table 1). Across diseases, the number of candidate drugs (Bonferroni-corrected p-value ≤ 0.05) generally increased across the three validation scenarios, with most diseases showing a stepwise increase. Notable exceptions included Alzheimer’s disease and lymphoma, which showed an increase in candidate drugs following network updating and a subsequent decrease after gene expansion, as well as breast cancer, which exhibited a decrease after network updating followed by an increase after gene expansion. Importantly, for all three exceptions, candidate drug counts with DGLinker-expanded gene sets remained higher than those observed under the old network.

Updating the network generally improved the recovery of known drugs (matching drug counts) in the DrugBank validation, with the exception of lymphoma and type 2 diabetes. However, this improvement was more limited in the MEDI2-HPS validation, where higher matching counts were only seen for Alzheimer’s disease and rheumatoid arthritis. When the Updated interaction network was held constant, replacing known disease genes with DGLinker-expanded gene sets resulted in heterogeneous changes in the number of approved drugs identified: in DrugBank, increases were limited to asthma and schizophrenia, while under MEDI2-HPS, slight increases were observed for asthma and hypertension; other diseases showed unchanged or reduced true positive counts.

#### Impact of ATCEA on Validation Metrics and Candidate Retention

To evaluate the contribution of ATCEA within our drug repurposing pipeline, we assessed precision, recall, F1-score, accuracy, and specificity across the twelve diseases in both DrugBank and MEDI2-HPS validation datasets before and after ATCEA filtering under the three validation scenarios (Figure 3A-B). The complete disease-specific ATCEA enrichment results are provided in Supplement Document 2, and the full validation metrics are provided in Supplement Documents 3 and 4.

**Figure 3.**
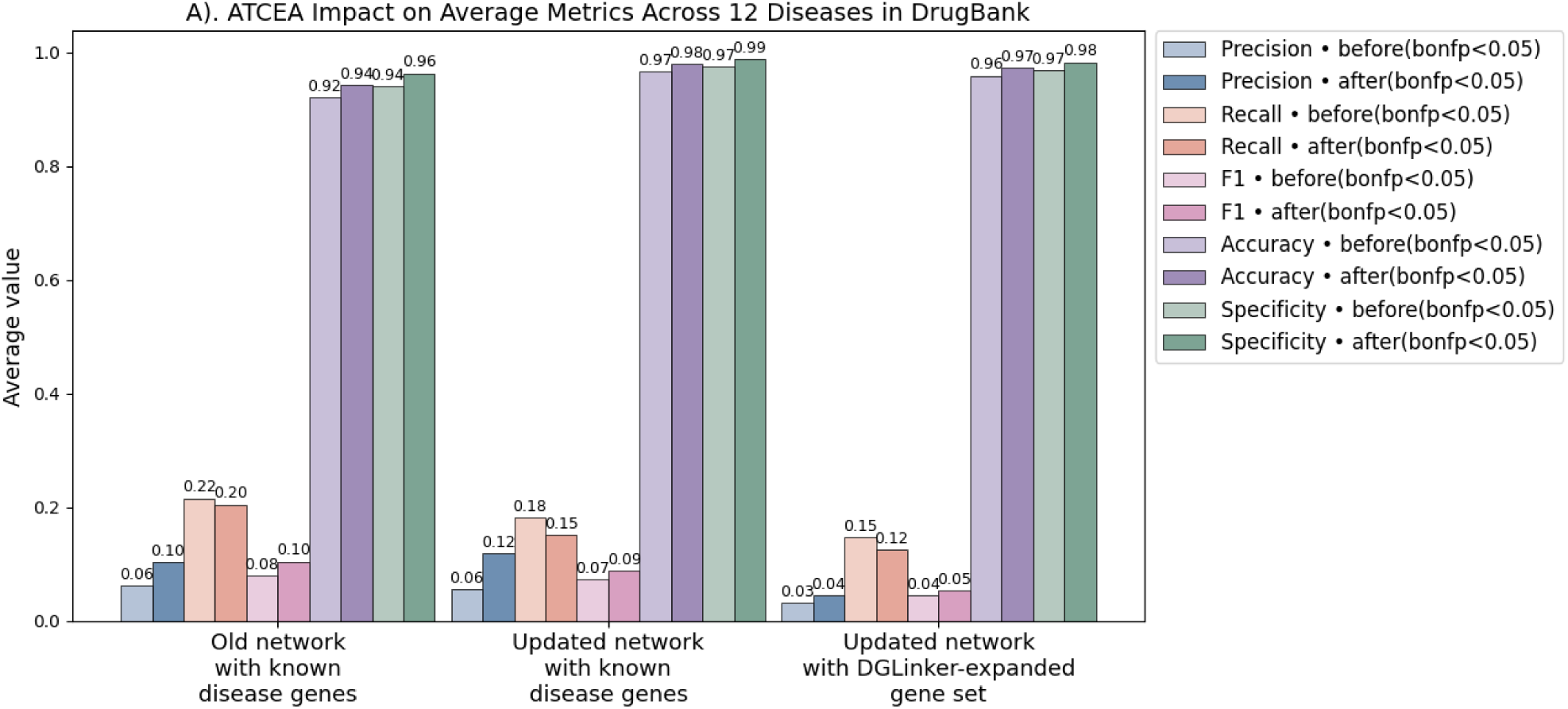
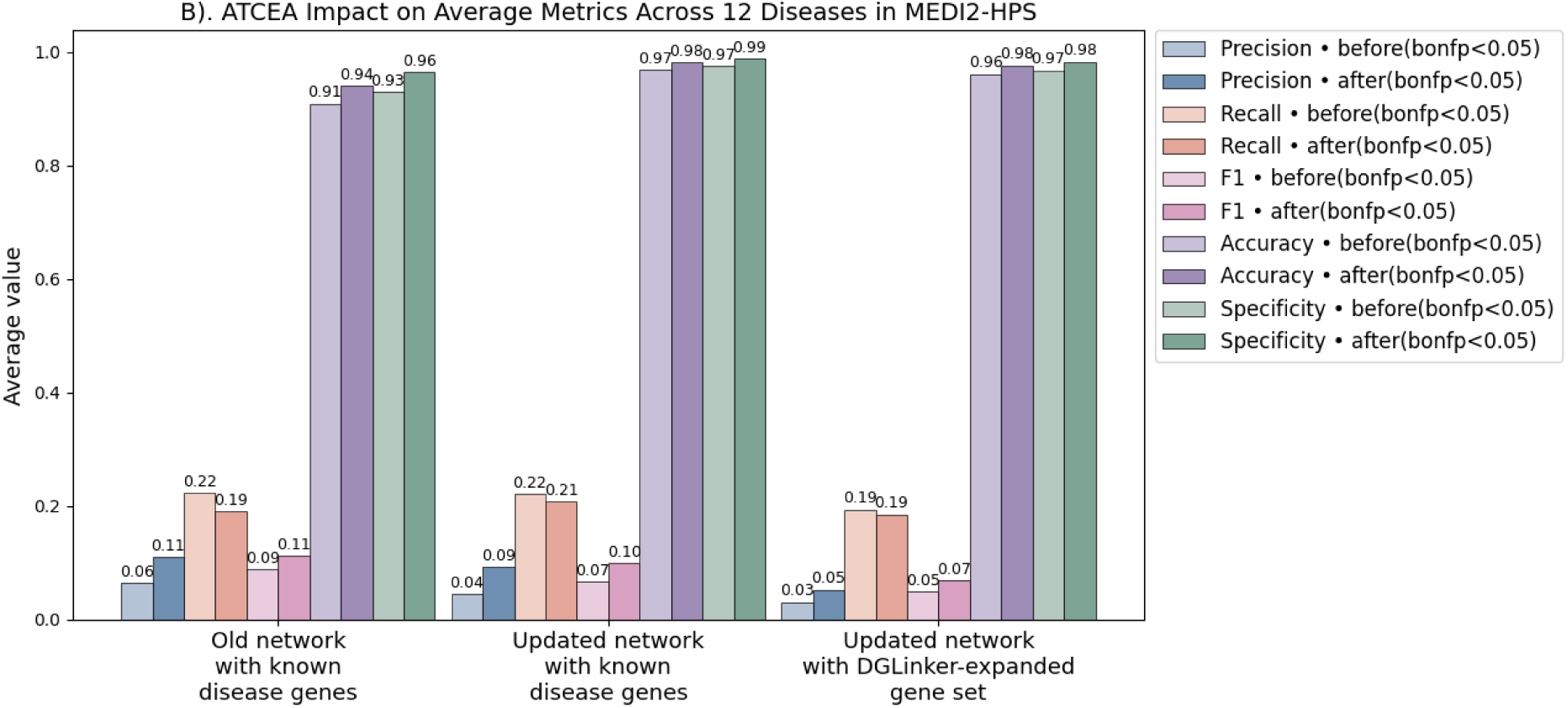
Average metrics comparison of drug repurposing results in different databases.

Across both validation databases, ATCEA filtering resulted in consistent increases in precision, F1-score, and decreases in recall (Figure 3A-B). In the DrugBank validation (Figure 3A), ATCEA increased average precision (e.g., from 0.06 to 0.12 in the updated network with known disease genes), together with corresponding improvements in F1-score and specificity, while recall showed a modest decrease (from 0.18 to 0.15). A similar directional trend was observed in the MEDI2-HPS validation (Figure 3B), where precision increased (from 0.04 to 0.09 under the same validation scenario), alongside improvements in F1-score and specificity, whereas recall slightly decreased (from 0.22 to 0.21).

These results indicate that ATCEA operates as a conservative post hoc filter that favours higher-confidence drug-disease associations over exhaustive coverage, a pattern that is further examined through candidate drugs retention analysis in Figure 4.

**Figure 4.**
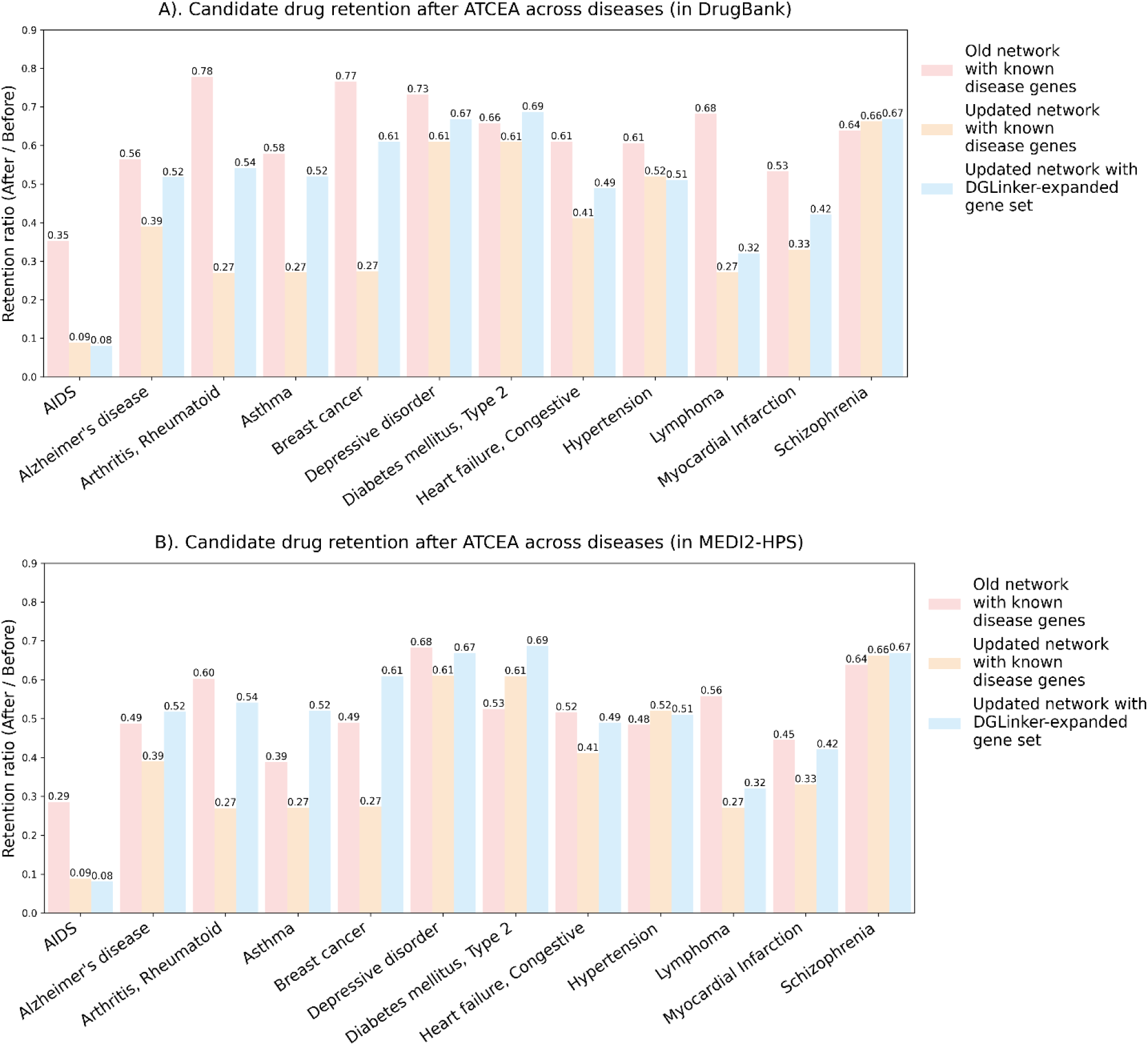
Disease-specific candidate drugs retention after ATCEA in different databases. Candidate drug retention is defined as the ratio of the number of candidate drugs retained after ATCEA to the number of candidate drugs before ATCEA, for each disease and validation scenario. Results are shown for DrugBank (A) and MEDI2-HPS (B).

We also quantified candidate drugs retention after ATCEA across diseases and network configurations (Figure 4A-B). ATCEA substantially reduced the number of retained candidate drugs, with retention ratios varying across diseases and network settings. Lower retention was generally observed in the updated and DGLinker-expanded networks, whereas several diseases consistently retained a larger fraction of candidates across validation scenarios.

These candidate retention results explain the reduction in recall observed after ATCEA in Figure 3. By restricting predictions to significantly enriched ATC categories, ATCEA reduces the size of the retained prediction set and removes some known drug-disease associations. Because the total number of known drugs remains fixed, removing TP from the retained set directly leads to a decrease in recall.

At the same time, although the total number of retained predictions (TP + FP) decreases, average precision increases slightly, indicating that false positives are removed more strongly than true positives. Together, these results reflect a trade-off in which ATCEA narrows the candidate drug space and improves prioritisation at the expense of coverage, rather than altering the underlying drug ranking.

In a subset of diseases, ATCEA led to particularly stringent filtering, such that no known drug-disease associations were retained after ATCEA (see Supplement Document 3 and 4).

### Drug Repurposing on ALS

Many human diseases have few or no effective treatments available, which makes the systematic evaluation of drug repurposing strategies challenging, despite these being the conditions that could benefit most from such approaches. ALS exemplifies this scenario. In the following case study, we apply our pipeline to ALS and validate the results through a manual assessment of the clinical relevance of the predicted drugs.

#### Disease-Gene Set Construction and Comparison

To investigate how knowledge-graph-assisted gene expansion influences downstream drug predictions and pharmacological enrichment, we curated three distinct ALS-associated gene sets (Figure 5): known ALS gene set (N = 104), new predicted ALS gene set from DGLinker (N = 972), and all ALS gene set which included both known and predicted ALS genes (N = 1076).

**Figure 5.**
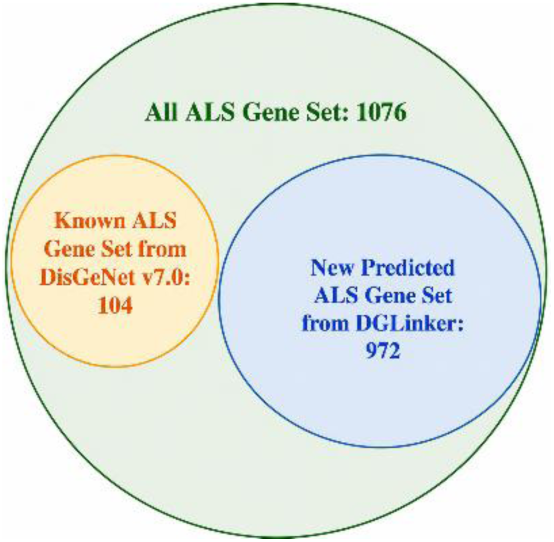
ALS gene set sizes: In yellow Known ALS gene set from DisGeNET (Piñero et al., 2020); In blue New predicted ALS gene set is from DGLinker (Hu et al., 2021), we leveraged DisGeNet v7.0 (Piñero et al., 2020), Gene Ontology v2021-02 (Consortium et al., 2023), and IntAct v2021-04 (del Toro et al., 2022) databases as input options; In green we combined the known ALS gene set with the new predicted ALS gene set as all ALS gene set, to create a more extensive database.

#### SAveRUNNER-Derived Candidate Drugs

SAveRUNNER was applied to each gene set to rank candidate drugs based on similarity. Following Bonferroni-corrected p-value<0.05, we identified 64 significant candidate drugs from the known ALS gene set, 128 from the DGLinker-new predicted ALS gene set, and 133 from all ALS gene set. Full results are in Table S1 in Supplement Document 1.

These results demonstrate that incorporating predicted genes by DGLinker doubles the number of significant drug associations compared to curated knowledge alone (from 64 to 133 candidate drugs), indicating a substantially broader repurposing landscape when novel predicted genes are included.

#### ATCEA and Candidate Drugs Prioritisation

To visualize the ATC enrichment analysis results, we constructed a heatmap to show the relationships between ATC categories and the three gene sets (Figure 6).

**Figure 6.**
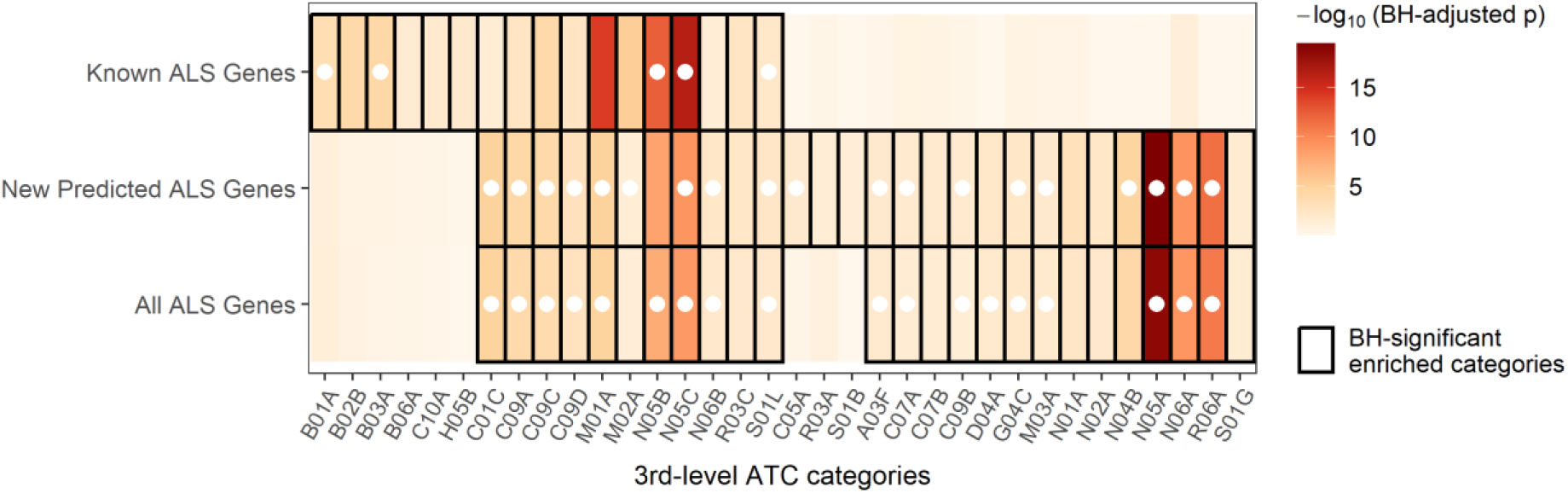
Heatmap: ATCEA results across different ALS gene sets. Color intensity indicating the –log10(BH-adjusted p-value) for each category. Cells outlined with a black border denote categories that are significantly enriched (BH-adjusted p < 0.05) within the corresponding gene set. White circles indicate that the ATC categories contain at least one Bonferroni-significant SAveRUNNER drug.

Using ATCEA on the complete SAveRUNNER drug-disease network, we initially identified 34 enriched third-level ATC categories across different ALS gene sets (Figure 6). We traced these enriched ATC categories back to the SAveRUNNER drug results and required that each category contained at least one Bonferroni-significant drug, only 23 ATC categories remained, which we regarded as the final set of candidate categories. Our analysis revealed 77 unique Bonferroni-significant drugs within BH-adjusted significant ATC categories (Table 2), including 22 drugs identified exclusively from analyses based on known ALS genes. We then assessed the impact of different gene sets on the ATCEA results, there were 11 ATC categories, including **N06A (antidepressants)** and **N05A (antipsychotics)**, that were only significantly enriched in the *New predicted ALS gene set* and *All ALS gene set*, but not in the *known ALS gene set* (Figure 6). Moreover, we observed that many drugs within these enriched ATC categories are supported by emerging evidence for ALS therapeutic relevance, derived from mechanistic studies, cellular and animal models, epidemiological observations, or independent computational repurposing analyses (Table 2). Notably, all candidate drugs with such supporting evidence were Bonferroni-significant only in the *New predicted ALS gene set* and *All ALS gene set*. This finding highlights the methodological value of incorporating DGLinker-derived gene expansion into our drug repurposing pipeline.

**Table 2.**
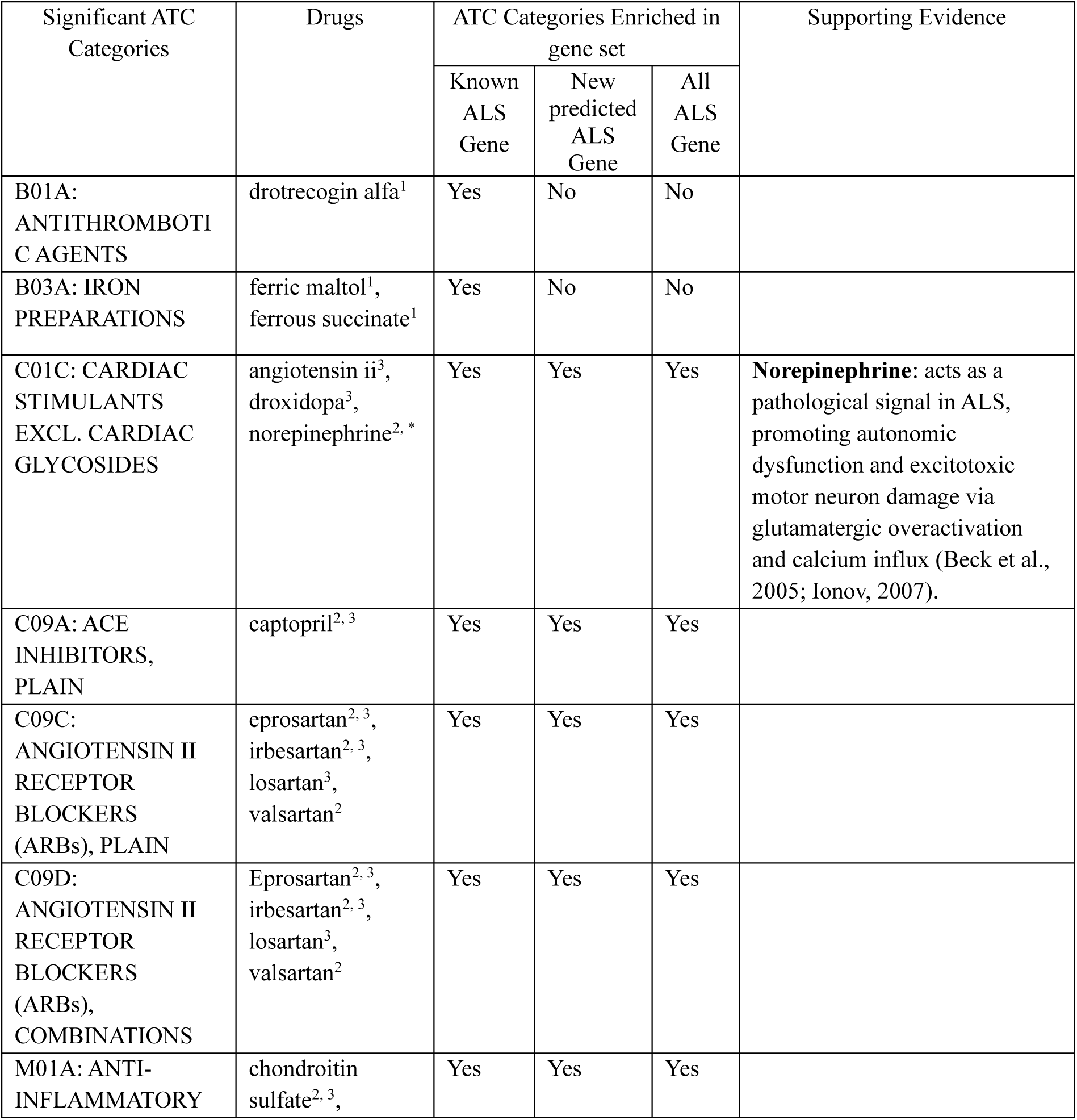
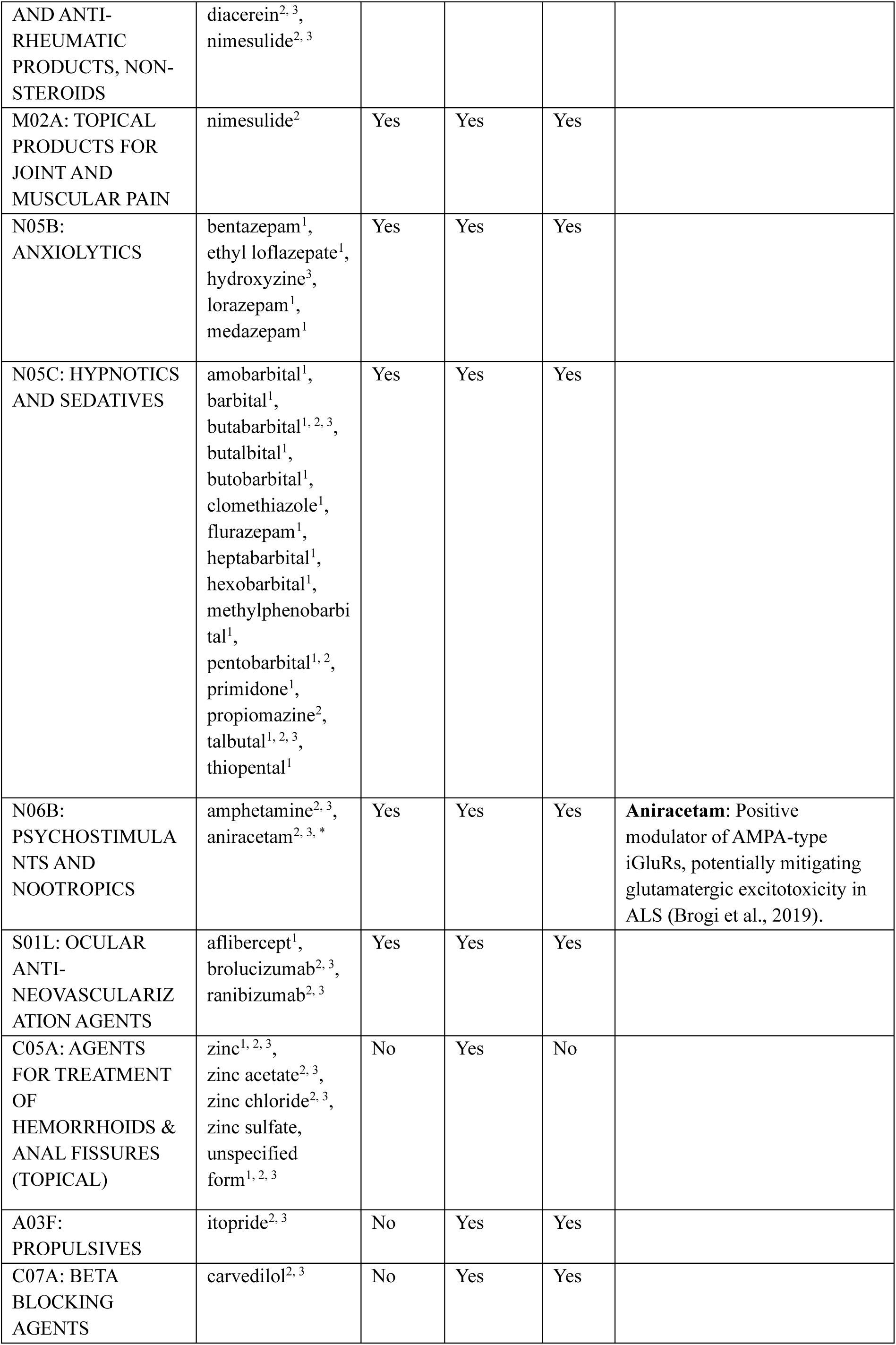
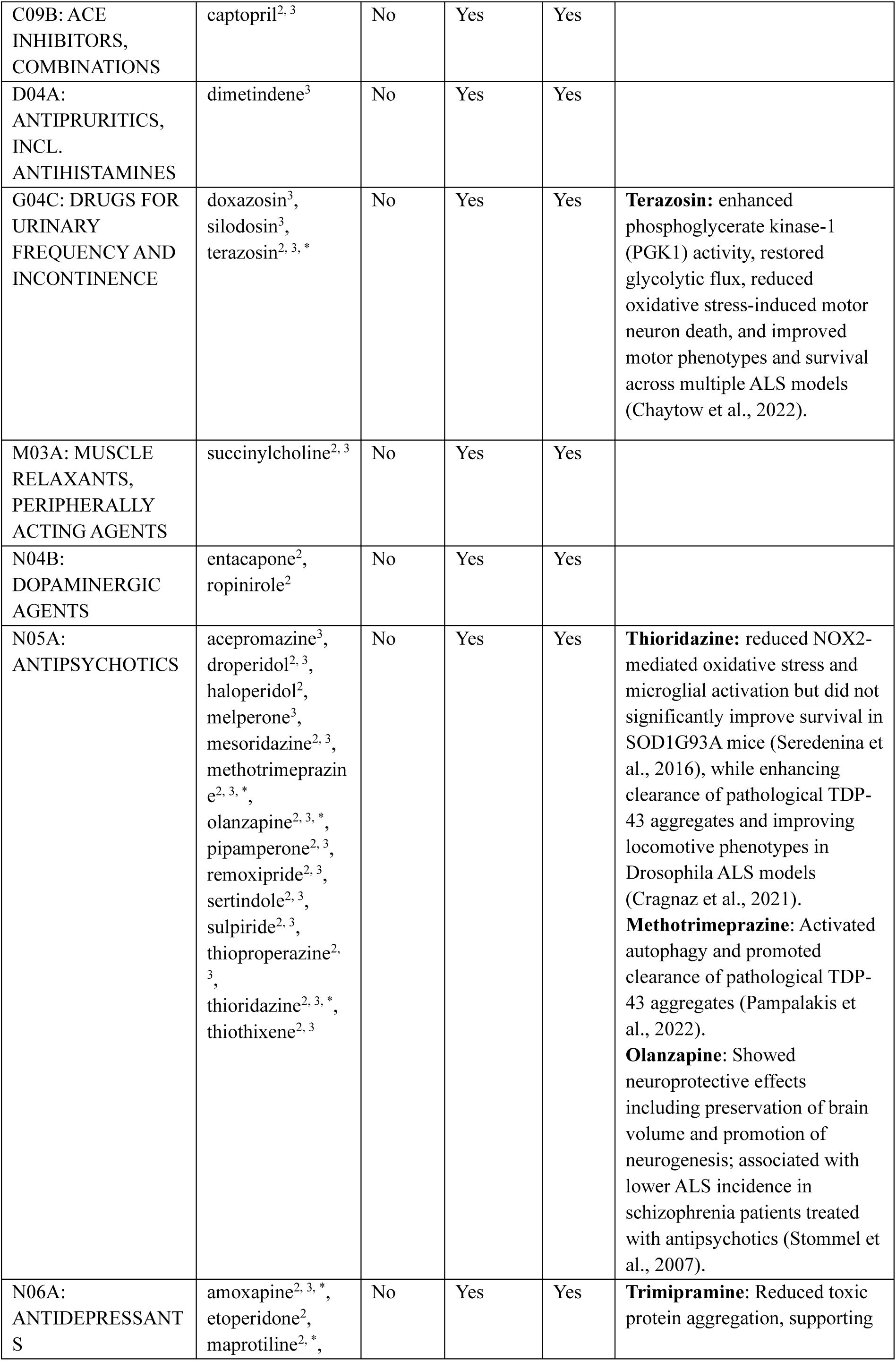
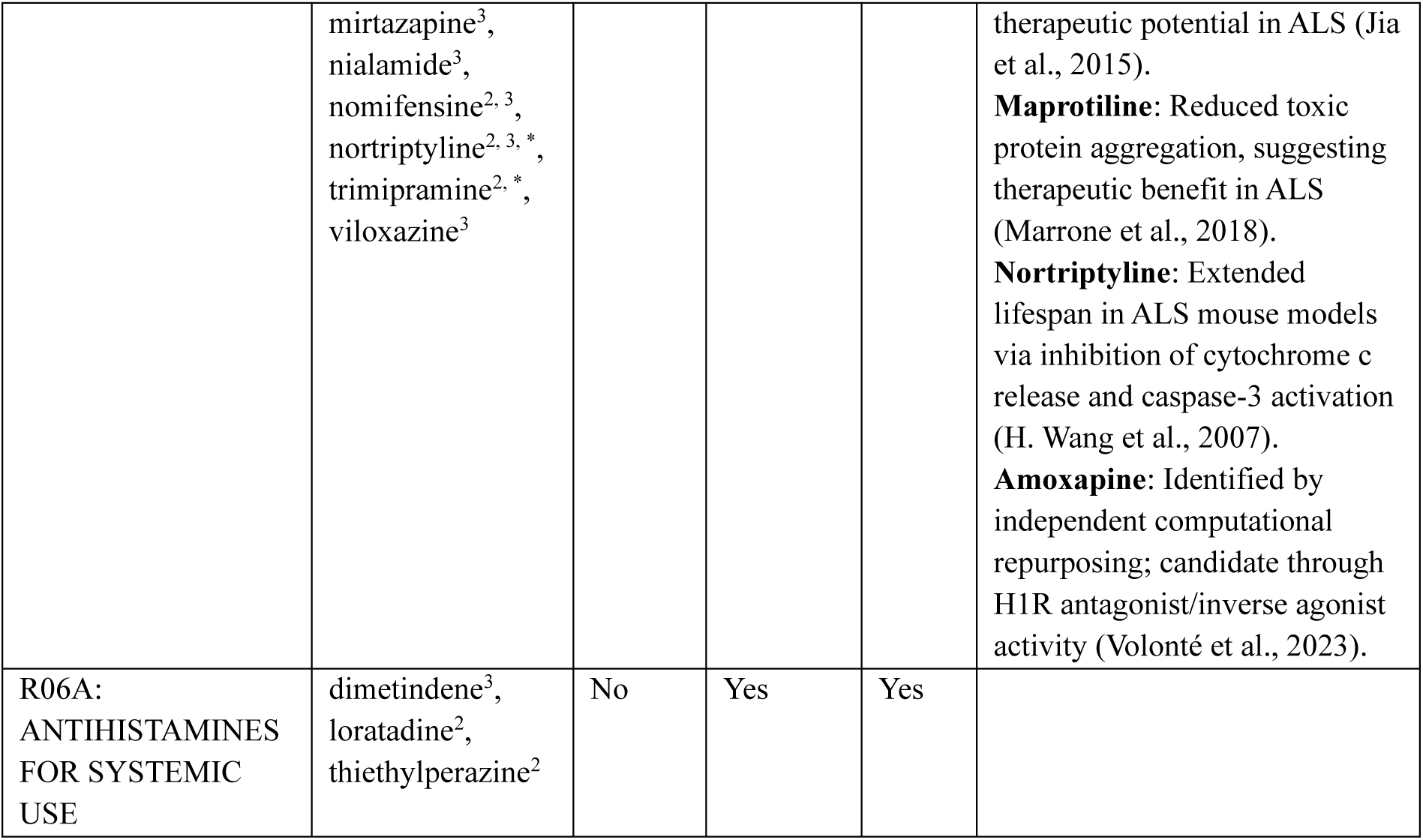
Candidate Drugs in the Significant 3^rd^ level ATC Categories. Drug with superscripts 1 if drug Bonferroni-significant in known ALS genes, 2 if Bonferroni-significant in new predicted ALS genes, 3 if Bonferroni-significant in all ALS genes. Drugs with “*” indicate that there is related published research on their investigation as potential treatment for ALS.

Several candidate drugs identified by our pipeline have evidence suggesting potential relevance to ALS. For example, **nortriptyline** has been reported to extend survival in ALS mouse models via anti-apoptotic pathways (H. Wang et al., 2007); **methotrimeprazine** has been shown to enhance autophagic clearance of TDP-43 aggregates, a pathological feature of ALS (Pampalakis et al., 2022); and **aniracetam**, an AMPA receptor modulator, has been proposed to mitigate glutamate-mediated excitotoxicity, a mechanism implicated in ALS progression (Brogi et al., 2019). This supports the potential utility of the pipeline in selecting compounds with mechanistic features consistent with ALS pathogenesis.

#### EHR Evaluation of ALS Candidate Drugs

Among the 77 prioritised ALS candidate drugs, 15 were identified in the EHR prescription records. Of these, 14 showed at least one pre-diagnosis exposure, 7 had sufficient data for pre-diagnosis survival analysis, and 13 showed at least one post-diagnosis exposure, of which 3 were evaluable in the post-diagnosis landmark analysis. None of the evaluated drugs remained significant after Benjamini-Hochberg correction.

In the pre-diagnosis analysis, doxazosin showed the lowest hazard ratio (HR = 0.516, 95% CI 0.244-1.095), suggesting longer survival relative to matched controls. Mirtazapine and irbesartan also showed HR below 1, whereas losartan showed a near-null association (HR = 0.899, 95% CI 0.450-1.798). In contrast, lorazepam (HR = 2.671, 95% CI 0.994-7.178) and haloperidol (HR = 5.426, 95% CI 1.034-28.475) showed HR above 1, corresponding to shorter survival in exposed patients relative to matched controls.

In the post-diagnosis landmark analysis, loratadine showed an HR of 1.166 (95% CI 0.22-6.10), indicating little apparent difference from matched controls, whereas lorazepam showed a higher HR (3.148, 95% CI 0.62-15.92). Mirtazapine showed an HR below 1 (0.039, 95% CI 0.00-1.93), although confidence intervals were wide for all three drugs, indicating substantial uncertainty due to the small sample size (Figure 7).

**Figure 7.**
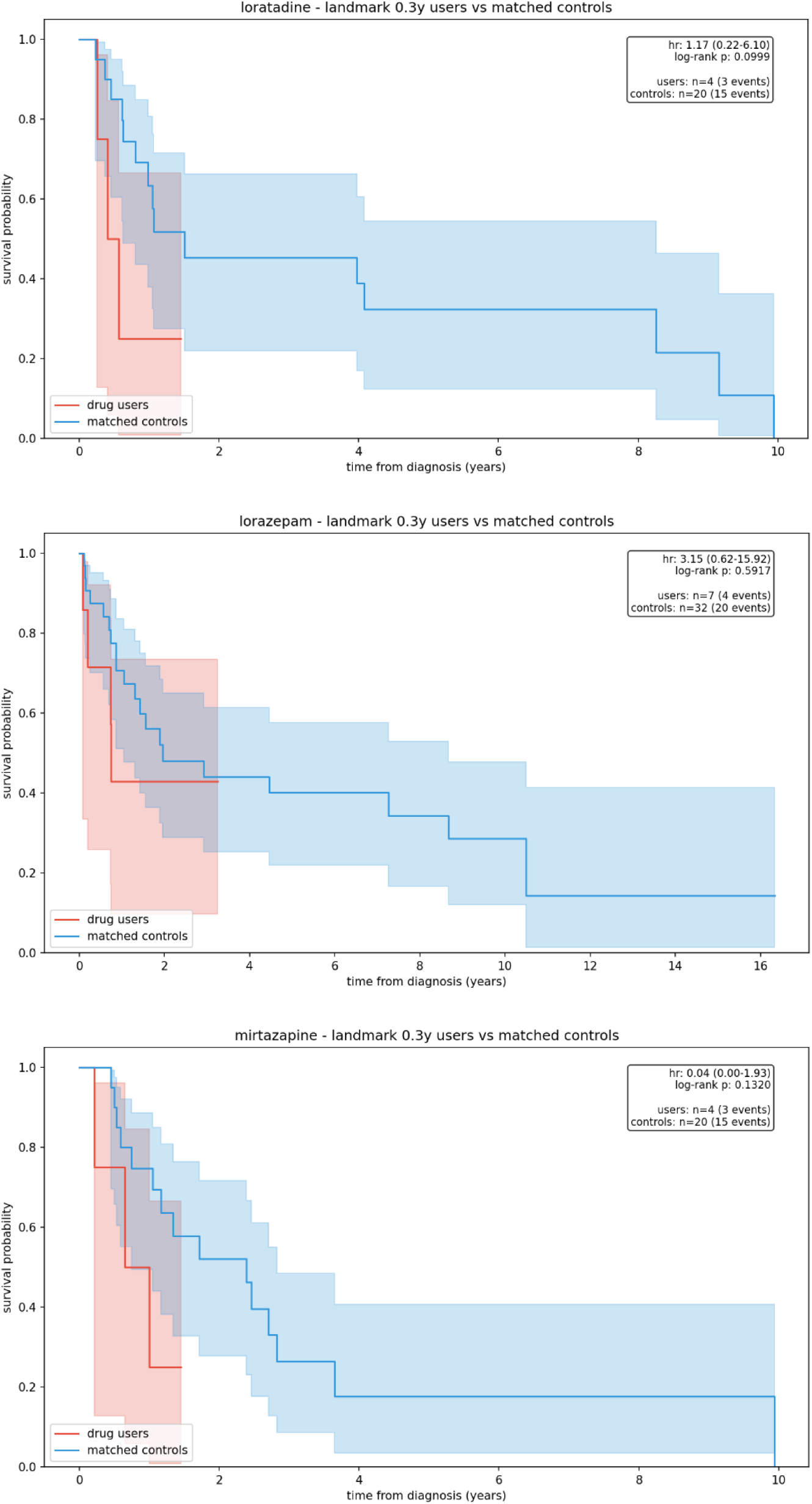
Post-diagnosis landmark Kaplan-Meier survival curves for evaluable ALS candidate drugs. HRs are shown relative to matched controls. HR < 1 indicates longer survival among exposed patients, whereas HR > 1 indicates shorter survival; values close to 1 suggest little difference between groups. Log-rank p-values test for differences in survival distributions; ’events’ indicate the number of observed deaths.

At the pharmacological class level, only the pre-diagnosis analysis yielded ATC categories represented by at least two drugs, which more meaningfully reflect class-level aggregation than categories represented by a single drug. Under this criterion, ARBs (C09C, drugs = 3, HR = 1.024), antidepressants (N06A, drugs = 2, HR = 0.247), and antipsychotics (N05A, drugs = 2, HR = 5.426) were evaluable. Antidepressants showed a HR below 1, antipsychotics showed a HR above 1, and ARBs showed a near-null effect. These pre-diagnosis class-level findings provide exploratory context for interpreting the ATCEA results. Full EHR evaluation results are provided in Supplement Document 5.

## Discussion

In this study, we present an integrative drug repurposing pipeline that combines computationally predicted gene-disease associations (DGLinker), network-based drug prioritisation (SAveRUNNER), and pharmacological class-level filtering through ATC Category Enrichment Analysis (ATCEA). Our analysis yielded three main findings. First, benchmark validation showed that updating the network and incorporating DGLinker-predicted genes generally increased the number of predicted candidate drugs across diseases, while AUROC values remained stable, indicating preserved discriminative performance across validation scenarios and databases. Second, applying ATCEA as a post hoc filter led to modest but directionally consistent increases in average precision, F1-score, and specificity, accompanied by reduced recall and contraction of the candidate drugs space. Third, when applied to amyotrophic lateral sclerosis (ALS), the pipeline identified 77 candidate drugs within significantly enriched pharmacological classes, several of which have independent support from preclinical studies, highlighting the biological relevance of the predictions. These findings are discussed in detail below.

The AUROC results indicate that the proposed drug repurposing pipeline provides a robust discrimination framework while flexibly adapting to updates in biological knowledge. Across diseases, updating the network and incorporating DGLinker-expanded gene sets resulted in only small, disease-specific changes in AUROC, suggesting that discriminative performance is preserved across validation scenarios and databases. This observation is consistent with the robustness of the network-based scoring framework implemented in SAveRUNNER.

In contrast, network updating and gene expansion primarily affected the size and composition of the candidate drugs space, leading to increased numbers of significant candidate drugs for most diseases, with clear disease-specific variation. Changes in the recovery of known drug-disease associations were heterogeneous and not consistently positive, particularly in DrugBank and MEDI2-HPS validations. These patterns likely reflect differences in validation resource coverage and annotation completeness. Indeed, it is important to highlight that the use of known drug-disease associations from DrugBank and MEDI2-HPS represents an incomplete set of true positives and an imperfect set of true negatives. Some drugs currently annotated as relevant to a given disease may also be relevant to other diseases, but such associations have not yet been identified or curated. Moreover, our drug predictions depend on the genetic definition of the target disease, which remains substantially limited for most human diseases. These are limiting factors in our validation strategy and should be considered.

Following network-based prediction and gene expansion, ATCEA was applied as a post hoc step to reorganize drug-level predictions into pharmacological classes defined by shared ATC annotations. By aggregating drugs at the class level, ATCEA highlights pharmacologically coherent signals rather than treating candidate drugs as independent entities. This class-based aggregation systematically narrows the candidate drugs space, resulting in a balance between coverage and precision. In drug repurposing settings characterized by large and heterogeneous candidate lists, this class-level view supports more focused interpretation by prioritising recurrent therapeutic categories among top-ranked candidates.

These validation results indicate stable discriminative performance, alongside scenario-specific variation in the composition of the candidate drug space. Taken together, the integration of disease gene expansion, network-based statistical association analysis, and systematic classification of candidate drugs defines a modular framework applicable to diverse disease contexts, particularly where therapeutic annotations are limited.

Application of the pipeline to ALS highlights the impact of disease gene expansion on candidate drug prioritisation in a setting where the target disease lacks enough known drugs to enable systematic evaluation of drug repurposing methods. When DGLinker-expanded gene sets were incorporated, ATC category enrichment revealed therapeutic classes that were not detected using known ALS genes alone, including antipsychotics (N05A), antidepressants (N06A), and other categories. This suggests that gene expansion may capture broader molecular processes linked to ALS beyond those represented by currently known disease genes. Some of the prioritised drugs within these newly identified categories have reported associations with biological mechanisms relevant to ALS pathophysiology. For example, nortriptyline has been linked to modulation of apoptotic pathways, while methotrimeprazine has been reported to influence clearance of TDP-43 aggregates. These observations provide biological context for the pipeline outputs and support its use as a hypothesis-generating framework.

The EHR evaluation added a further exploratory dimension to this ALS case study by providing clinical follow-up for a subset of the computationally prioritised candidates. However, only a limited proportion of the prioritised ALS drugs could be followed up in the EHR, and even fewer had sufficient exposure for survival analysis once treatment timing was considered. As a result, the findings at the individual drug level should not be interpreted as validation of specific therapeutic effects, but rather as preliminary clinical context for a limited subset of prioritised candidates in a real-world ALS cohort.

At the pharmacological class level, the EHR analysis provided only limited additional context for the ATCEA findings. This limited interpretability also differed by exposure timing. The post-diagnosis analysis is conceptually more relevant to the therapeutic repurposing question, whereas the pre-diagnosis results require more cautious interpretation because they may reflect prodromal features, comorbidity patterns, or indication-related prescribing rather than direct treatment effects. In our study, interpretable class-level patterns were observed only in the pre-diagnosis analysis. A recent population-based study reported that pre-diagnosis use of common psychiatric medications was associated with both a higher subsequent risk of ALS and poorer prognosis after diagnosis (Chourpiliadis et al., 2025), supporting a cautious interpretation of the psychiatric-related ATC signals identified in our ALS case study. Taken together, the EHR findings are best interpreted as supportive evidence that does not establish causality, complementing the computational prioritisation framework rather than confirming therapeutic benefit. This framework may provide a useful basis for future EHR-based evaluation of computationally prioritised drug candidates in ALS.

Despite the strengths of our drug repurposing pipeline, several methodological limitations should be acknowledged. In the current implementation, disease-associated genes are treated as an unweighted set within SAveRUNNER, without accounting for gene-level relevance or confidence. As a result, the integration of DGLinker-predicted genes does not consistently increase the recovery of known drug-disease associations, as predicted genes that are less central to disease mechanisms may contribute limited additional signal to proximity-based scoring. Future extensions of the pipeline could address this limitation by incorporating gene importance weights, for example by prioritising curated disease genes or integrating DGLinker prediction scores into the proximity calculation.

Another notable limitation of this study is the lack of consideration for the direction of gene-disease and drug-gene effects. Consequently, it remains unclear whether the candidate drugs would exert beneficial or harmful outcomes. These findings should therefore be interpreted in conjunction with further experimental or clinical validation. Relatedly, the substantial variability in pipeline performance observed across different diseases suggests that the framework may be more effective for certain disease types than others.

In this context, an additional limitation concerns the application of ATC enrichment analysis as a post hoc filter. For certain diseases, restricting candidates to significantly enriched ATC categories resulted in zero true positives, reflecting the stringency of class-level aggregation. While this approach enhances interpretability and precision in some settings, it may be overly restrictive for diseases with sparse or heterogeneous therapeutic landscapes. Accordingly, the suitability of ATCEA, and the disease contexts in which it is most effective, warrant further investigation.

Future extensions of the pipeline could incorporate complementary sources of evidence. Subsequent research should also focus on evaluating alternative components beyond DGLinker and SAveRUNNER, further validating predictive performance through in silico analyses, and exploring applications to additional disease subtypes. These efforts will be essential for assessing disease-specific performance and establishing the broader utility of the framework across heterogeneous biological contexts.

In conclusion, this study demonstrates that integrating disease-gene prediction, network-based drug identification, and pharmacological class-level prioritisation provides a scalable and interpretable framework for drug repurposing, particularly for diseases with limited therapeutic options and incomplete genetic characterization. By integrating computational gene discovery with drug-level and class-level prioritisation, the proposed pipeline expands the therapeutic candidate space while preserving biological and pharmacological interpretability. Its application to ALS identified candidate drugs with biological and clinical relevance, supported by literature evidence and exploratory EHR evaluation, underscoring its value as a practical framework for prioritising candidates for subsequent experimental and clinical investigation.

## Acknowledgements

The authors acknowledge use of the King’s Computational Research, Engineering and Technology Environment (CREATE) (https://create.kcl.ac.uk), which is delivered in partnership with the National Institute for Health and Care Research (NIHR) Biomedical Research Centres at South London and Maudsley and Guy’s and St. Thomas’ NHS Foundation Trusts and part-funded by capital equipment grants from the Maudsley Charity (award 980) and Guy’s and St. Thomas’ Charity (TR130505).

## Ethics approval and data governance

This project was conducted in accordance with the ethical principles outlined in the Declaration of Helsinki and complied with all applicable data protection and governance regulations for the use of EHR data. The study used routinely collected, de-identified clinical data accessed within secure research environments, with appropriate safeguards to protect patient confidentiality and privacy. Data access and analysis were approved through the King’s Electronic Records Research Interface (KERRI).

## Reference

Al-Chalabi, A., Fang, F., Hanby, M. F., Leigh, P. N., Shaw, C. E., Ye, W., & Rijsdijk, F. (2010). An estimate of amyotrophic lateral sclerosis heritability using twin data. *Journal of Neurology*, Neurosurgery & Psychiatry, 81(12), 1324–1326. 10.1136/JNNP.2010.207464

Atassi, N., Cook, A., Pineda, C. M. E., Yerramilli-Rao, P., Pulley, D., & Cudkowicz, M. (2010). Depression in amyotrophic lateral sclerosis. Amyotrophic Lateral Sclerosis : Official Publication of the World Federation of Neurology Research Group on Motor Neuron Diseases, 12(2), 109. 10.3109/17482968.2010.536839

Bean, D. M., Al-Chalabi, A., Dobson, R. J. B., & Iacoangeli, A. (2020). A Knowledge-Based Machine Learning Approach to Gene Prioritisation in Amyotrophic Lateral Sclerosis. Genes 2020, Vol. 11, Page 668, 11(6), 668. 10.3390/GENES11060668

Beck, M., Flachenecker, P., Magnus, T., Giess, R., Reiners, K., Toyka, K. V., & Naumann, M. (2005). Autonomic dysfunction in ALS: A preliminary study on the effects of intrathecal BDNF. Amyotrophic Lateral Sclerosis, 6(2), 100–103. 10.1080/14660820510028412

Brogi, S., Campiani, G., Brindisi, M., & Butini, S. (2019). Allosteric Modulation of Ionotropic Glutamate Receptors: An Outlook on New Therapeutic Approaches to Treat Central Nervous System Disorders. ACS Medicinal Chemistry Letters, 10(3), 228–236. 10.1021/ACSMEDCHEMLETT.8B00450

Brunetti, M., Paci, P., & Fiscon, G. (2023). A network-based bioinformatic analysis for identifying potential repurposable active molecules in different types of human cancers. Proceedings - 2023 2023 IEEE International Conference on Bioinformatics and Biomedicine, BIBM 2023, 3626–3631. 10.1109/BIBM58861.2023.10385812

Cai, L., Chu, J., Xu, J., Meng, Y., Lu, C., Tang, X., Wang, G., Tian, G., & Yang, J. (2023). Machine learning for drug repositioning: Recent advances and challenges. Current Research in Chemical Biology, 3, 100042. 10.1016/J.CRCHBI.2023.100042

Cai, L., Lu, C., Xu, J., Meng, Y., Wang, P., Fu, X., Zeng, X., & Su, Y. (2021). Drug repositioning based on the heterogeneous information fusion graph convolutional network. Briefings in Bioinformatics, 22(6), 1–12. 10.1093/BIB/BBAB319

Chaytow, H., Carroll, E., Gordon, D., Huang, Y. T., van der Hoorn, D., Smith, H. L., Becker, T., Becker, C. G., Faller, K. M. E., Talbot, K., & Gillingwater, T. H. (2022). Targeting phosphoglycerate kinase 1 with terazosin improves motor neuron phenotypes in multiple models of amyotrophic lateral sclerosis. EBioMedicine, 83, 104202. 10.1016/j.ebiom.2022.104202

Cheng, F., Desai, R. J., Handy, D. E., Wang, R., Schneeweiss, S., Barabási, A. L., & Loscalzo, J. (2018). Network-based approach to prediction and population-based validation of in silico drug repurposing. Nature Communications 2018 9:1, 9(1), 1–12. 10.1038/s41467-018-05116-5

Chourpiliadis, C., Lovik, A., Ingre, C., Press, R., Samuelsson, K., Valdimarsdottir, U., & Fang, F. (2025). Use of Common Psychiatric Medications and Risk and Prognosis of Amyotrophic Lateral Sclerosis. JAMA Network Open, 8(6), e2514437–e2514437. 10.1001/JAMANETWORKOPEN.2025.14437

Consortium, T. G. O., Aleksander, S. A., Balhoff, J., Carbon, S., Cherry, J. M., Drabkin, H. J., Ebert, D., Feuermann, M., Gaudet, P., Harris, N. L., Hill, D. P., Lee, R., Mi, H., Moxon, S., Mungall, C. J., Muruganugan, A., Mushayahama, T., Sternberg, P. W., Thomas, P. D., … Westerfield, M. (2023). The Gene Ontology knowledgebase in 2023. Genetics, 224(1). 10.1093/GENETICS/IYAD031

Cook, C., & Petrucelli, L. (2019). Genetic Convergence Brings Clarity to the Enigmatic Red Line in ALS. Neuron, 101(6), 1057–1069. 10.1016/J.NEURON.2019.02.032

Cragnaz, L., Spinelli, G., De Conti, L., Bureau, E. A., Brownlees, J., Feiguin, F., Romano, V., Skoko, N., Klima, R., Kettleborough, C. A., Baralle, F. E., & Baralle, M. (2021). Thioridazine reverts the phenotype in cellular and Drosophila models of amyotrophic lateral sclerosis by enhancing TDP-43 aggregate clearance. Neurobiology of Disease, 160, 105515. 10.1016/J.NBD.2021.105515

Davis, A. P., Wiegers, T. C., Sciaky, D., Barkalow, F., Strong, M., Wyatt, B., Wiegers, J., McMorran, R., Abrar, S., & Mattingly, C. J. (2025). Comparative Toxicogenomics Database’s 20th anniversary: update 2025. Nucleic Acids Research, 53(D1), D1328–D1334. 10.1093/NAR/GKAE883

del Toro, N., Shrivastava, A., Ragueneau, E., Meldal, B., Combe, C., Barrera, E., Perfetto, L., How, K., Ratan, P., Shirodkar, G., Lu, O., Mészáros, B., Watkins, X., Pundir, S., Licata, L., Iannuccelli, M., Pellegrini, M., Martin, M. J., Panni, S., … Hermjakob, H. (2022). The IntAct database: efficient access to fine-grained molecular interaction data. Nucleic Acids Research, 50(D1), D648–D653. 10.1093/NAR/GKAB1006

Fiscon, G., & Paci, P. (2021). SAveRUNNER: an R-based tool for drug repurposing. BMC Bioinformatics, 22(1), 1–10. 10.1186/S12859-021-04076-W

Fiscon, G., Sibilio, P., Funari, A., Conte, F., & Paci, P. (2022). Identification of Potential Repurposable Drugs in Alzheimer’s Disease Exploiting a Bioinformatics Analysis. Journal of Personalized Medicine, 12(10), 1731. 10.3390/JPM12101731/S1

Gallagher, G. W., Nowacek, D., Gutgsell, O., & Callaghan, B. C. (2021). Comparison of the United Kingdom and United States approaches to approval of new neuromuscular therapies. Muscle & Nerve, 64(6), 641–650. 10.1002/MUS.27380

Gleiss, A., Oberbauer, R., & Heinze, G. (2018). An unjustified benefit: immortal time bias in the analysis of time-dependent events. Transplant International, 31(2), 125–130. 10.1111/tri.13081

Hamad, A. A., Alkhawaldeh, I. M., Nashwan, A. J., Meshref, M., & Imam, Y. (2025). Tofersen for SOD1 amyotrophic lateral sclerosis: a systematic review and meta-analysis. Neurological Sciences 2025 46:5, 46(5), 1977–1985. 10.1007/S10072-025-07994-2

Hinchcliffe, M., & Smith, A. (2017). Riluzole: real-world evidence supports significant extension of median survival times in patients with amyotrophic lateral sclerosis. *Degenerative Neurological and Neuromuscular Disease*, Volume 7, 61–70. 10.2147/DNND.S135748

Hu, J., Lepore, R., Dobson, R. J. B., Al-Chalabi, A., Bean, D. M., & Iacoangeli, A. (2021). DGLinker: flexible knowledge-graph prediction of disease–gene associations. Nucleic Acids Research, 49(W1), W153–W161. 10.1093/NAR/GKAB449

Iacoangeli, A., Dilliott, A. A., Khleifat, A. Al, Andersen, P. M., Başak, N. A., Cooper-Knock, J., Corcia, P., Couratier, P., deCarvalho, M., Drory, V. E., Glass, J. D., Gotkine, M., Lerner, Y. M., Hardiman, O., Landers, J. E., McLaughlin, R. L., Pardina, J. S. M., Morrison, K., Pinto, S., … Farhan, S. M. K. (2025). Oligogenic structure of amyotrophic lateral sclerosis has genetic testing, counselling and therapeutic implications. *Journal of Neurology*, Neurosurgery & Psychiatry, 0(11), jnnp-2024-335364. 10.1136/JNNP-2024-335364

Ionov, I. D. (2007). Survey of ALS-associated factors potentially promoting Ca2+ overload of motor neurons. Amyotrophic Lateral Sclerosis, 8(5), 260–265. 10.1080/17482960701523124

Jia, J., & Le, W. (2015). Molecular network of neuronal autophagy in the pathophysiology and treatment of depression. Neuroscience Bulletin, 31(4), 427–434. 10.1007/S12264-015-1548-2/METRICS

Kim, Y.;, Cho, Y.-R., Kim, Y., & Cho, Y.-R. (2023). Predicting Drug–Gene–Disease Associations by Tensor Decomposition for Network-Based Computational Drug Repositioning. Biomedicines 2023, Vol. 11, Page 1998, 11(7), 1998. 10.3390/BIOMEDICINES11071998

Knox, C., Wilson, M., Klinger, C. M., Franklin, M., Oler, E., Wilson, A., Pon, A., Cox, J., Chin, N. E. L., Strawbridge, S. A., Garcia-Patino, M., Kruger, R., Sivakumaran, A., Sanford, S., Doshi, R., Khetarpal, N., Fatokun, O., Doucet, D., Zubkowski, A., … Wishart, D. S. (2024). DrugBank 6.0: the DrugBank Knowledgebase for 2024. Nucleic Acids Research, 52(D1), D1265–D1275. 10.1093/NAR/GKAD976

Kraljevic, Z., Searle, T., Shek, A., Roguski, L., Noor, K., Bean, D., Mascio, A., Zhu, L., Folarin, A. A., Roberts, A., Bendayan, R., Richardson, M. P., Stewart, R., Shah, A. D., Wong, W. K., Ibrahim, Z., Teo, J. T., & Dobson, R. J. B. (2021). Multi-domain clinical natural language processing with MedCAT: The Medical Concept Annotation Toolkit. Artificial Intelligence in Medicine, 117. 10.1016/j.artmed.2021.102083

Marrone, L., Poser, I., Casci, I., Japtok, J., Reinhardt, P., Janosch, A., Andree, C., Lee, H. O., Moebius, C., Koerner, E., Reinhardt, L., Cicardi, M. E., Hackmann, K., Klink, B., Poletti, A., Alberti, S., Bickle, M., Hermann, A., Pandey, U. B., … Sterneckert, J. L. (2018). Isogenic FUS-eGFP iPSC Reporter Lines Enable Quantification of FUS Stress Granule Pathology that Is Rescued by Drugs Inducing Autophagy. Stem Cell Reports, 10(2), 375–389. 10.1016/j.stemcr.2017.12.018

Mehta, P. R., Iacoangeli, A., Opie-Martin, S., Van Vugt, J. J. F. A., Al Khleifat, A., Bredin, A., Ossher, L., Andersen, P. M., Hardiman, O., Mehta, A. R., Fratta, P., Talbot, K., Başak, N. A., Corcia, P., Couratier, P., De Carvalho, M., Drory, V., Glass, J. D., Gotkine, M., … Al-Chalabi, A. (2022). The impact of age on genetic testing decisions in amyotrophic lateral sclerosis. Brain, 145(12), 4440–4447. 10.1093/BRAIN/AWAC279

Meng, Y., Lu, C., Jin, M., Xu, J., Zeng, X., & Yang, J. (2022). A weighted bilinear neural collaborative filtering approach for drug repositioning. Briefings in Bioinformatics, 23(2), 1–13. 10.1093/BIB/BBAB581

Nascimento, A. C. A., Prudêncio, R. B. C., & Costa, I. G. (2016). A multiple kernel learning algorithm for drug-target interaction prediction. BMC Bioinformatics, 17(1), 1–16. 10.1186/S12859-016-0890-3

Nelson, M. R., Tipney, H., Painter, J. L., Shen, J., Nicoletti, P., Shen, Y., Floratos, A., Sham, P. C., Li, M. J., Wang, J., Cardon, L. R., Whittaker, J. C., & Sanseau, P. (2015). The support of human genetic evidence for approved drug indications. Nature Genetics 2015 47:8, 47(8), 856–860. 10.1038/ng.3314

Novac, N. (2013). Challenges and opportunities of drug repositioning. Trends in Pharmacological Sciences, 34(5), 267–272. 10.1016/J.TIPS.2013.03.004

Ostell, J. M. (2012). Entrez: The NCBI Search and Discovery Engine. Lecture Notes in Computer Science (Including Subseries Lecture Notes in Artificial Intelligence and Lecture Notes in Bioinformatics), 7348 *LNBI*, 1–4. 10.1007/978-3-642-31040-9_1

Pampalakis, G., Angelis, G., Zingkou, E., Vekrellis, K., & Sotiropoulou, G. (2022). A chemogenomic approach is required for effective treatment of amyotrophic lateral sclerosis. Clinical and Translational Medicine, 12(1), e657. 10.1002/CTM2.657

Piñero, J., Ramírez-Anguita, J. M., Saüch-Pitarch, J., Ronzano, F., Centeno, E., Sanz, F., & Furlong, L. I. (2020). The DisGeNET knowledge platform for disease genomics: 2019 update. Nucleic Acids Research, 48(D1), D845–D855. 10.1093/NAR/GKZ1021

Reay, W. R., & Cairns, M. J. (2021). Advancing the use of genome-wide association studies for drug repurposing. Nature Reviews Genetics 2021 22:10, 22(10), 658–671. 10.1038/s41576-021-00387-z

Sadeghi, S., Lu, J., & Ngom, A. (2022). An Integrative Heterogeneous Graph Neural Network–Based Method for Multi-Labeled Drug Repurposing. Frontiers in Pharmacology, 13, 908549. 10.3389/FPHAR.2022.908549

Seredenina, T., Nayernia, Z., Sorce, S., Maghzal, G. J., Filippova, A., Ling, S. C., Basset, O., Plastre, O., Daali, Y., Rushing, E. J., Giordana, M. T., Cleveland, D. W., Aguzzi, A., Stocker, R., Krause, K. H., & Jaquet, V. (2016). Evaluation of NADPH oxidases as drug targets in a mouse model of familial amyotrophic lateral sclerosis. Free Radical Biology and Medicine, 97, 95–108. 10.1016/J.FREERADBIOMED.2016.05.016

Shah, S., Dooms, M. M., Amaral-Garcia, S., & Igoillo-Esteve, M. (2021). Current Drug Repurposing Strategies for Rare Neurodegenerative Disorders. Frontiers in Pharmacology, 12, 768023. 10.3389/FPHAR.2021.768023

Sibilio, P., Bini, S., Fiscon, G., Sponziello, M., Conte, F., Pecce, V., Durante, C., Paci, P., Falcone, R., Norata, G. D., Farina, L., & Verrienti, A. (2021). In silico drug repurposing in COVID-19: A network-based analysis. Biomedicine & Pharmacotherapy, 142, 111954. 10.1016/J.BIOPHA.2021.111954

Spargo, T. P., Gilchrist, L., Hunt, G. P., Dobson, R. J. B., Proitsi, P., Al-Chalabi, A., Pain, O., & Iacoangeli, A. (2024). Statistical examination of shared loci in neuropsychiatric diseases using genome-wide association study summary statistics. ELife, 12. 10.7554/ELIFE.88768

Stommel, E. W., Graber, D., Montanye, J., Cohen, J. A., & Harris, B. T. (2007). Does treating schizophrenia reduce the chances of developing amyotrophic lateral sclerosis? Medical Hypotheses, 69(5), 1021–1028. 10.1016/J.MEHY.2007.02.041

Sun, X., Jia, X., Lu, Z., Tang, J., & Li, M. (2024). Drug repositioning with adaptive graph convolutional networks. Bioinformatics, 40(1). 10.1093/BIOINFORMATICS/BTAD748

Van Daele, S. H., Moisse, M., Van Vugt, J. J. F. A., Zwamborn, R. A. J., Van Der Spek, R., Van Rheenen, W., Van Eijk, K., Kenna, K., Corcia, P., Vourc’h, P., Couratier, P., Hardiman, O., Mclaughin, R., Gotkine, M., Drory, V., Ticozzi, N., Silani, V., Ratti, A., De Carvalho, M., … Van Damme, P. (2023). Genetic variability in sporadic amyotrophic lateral sclerosis. Brain, 146(9), 3760–3769. 10.1093/BRAIN/AWAD120

Volonté, C., & Amadio, S. (2023). Amyotrophic lateral sclerosis disease burden: doing better at getting better. Neural Regeneration Research, 18(8), 0. 10.4103/1673-5374.363193

Wang, H., Guan, Y., Wang, X., Smith, K., Cormier, K., Zhu, S., Stavrovskaya, I. G., Huo, C., Ferrante, R. J., Kristal, B. S., & Friedlander, R. M. (2007). Nortriptyline delays disease onset in models of chronic neurodegeneration. European Journal of Neuroscience, 26(3), 633–641. 10.1111/J.1460-9568.2007.05663.X

Wang, M., Tang, C., & Chen, J. (2018). Drug-Target Interaction Prediction via Dual Laplacian Graph Regularized Matrix Completion. BioMed Research International, 2018(1), 1425608. 10.1155/2018/1425608

Wang, X., Blanchard, J., Grundke-Iqbal, I., Wegiel, J., Deng, H. X., Siddique, T., & Iqbal, K. (2014). Alzheimer disease and amyotrophic lateral sclerosis: An etiopathogenic connection. Acta Neuropathologica, 127(2), 243–256. 10.1007/S00401-013-1175-9

Wei, W. Q., Cronin, R. M., Xu, H., Lasko, T. A., Bastarache, L., & Denny, J. C. (2013). Development and evaluation of an ensemble resource linking medications to their indications. Journal of the American Medical Informatics Association, 20(5), 954–961. 10.1136/AMIAJNL-2012-001431

Xu, J., Mao, C., Hou, Y., Luo, Y., Binder, J. L., Zhou, Y., Bekris, L. M., Shin, J., Hu, M., Wang, F., Eng, C., Oprea, T. I., Flanagan, M. E., Pieper, A. A., Cummings, J., Leverenz, J. B., & Cheng, F. (2022). Interpretable deep learning translation of GWAS and multi-omics findings to identify pathobiology and drug repurposing in Alzheimer’s disease. Cell Reports, 41(9), 111717. 10.1016/J.CELREP.2022.111717

Yan, V. K. C., Li, X., Ye, X., Ou, M., Luo, R., Zhang, Q., Tang, B., Cowling, B. J., Hung, I., Siu, C. W., Wong, I. C. K., Cheng, R. C. K., & Chan, E. W. (2021). Drug Repurposing for the Treatment of COVID-19: A Knowledge Graph Approach. Advanced Therapeutics, 4(7), 2100055. 10.1002/ADTP.202100055

You, J., Islam, M. M., Grenier, L., Kuang, Q., McLeod, R. D., & Hu, P. (2018). Drug-target interaction network predictions for drug repurposing using LASSO-based regularized linear classification model. Lecture Notes in Computer Science (Including Subseries Lecture Notes in Artificial Intelligence and Lecture Notes in Bioinformatics), 10832 *LNAI*, 272–278. 10.1007/978-3-319-89656-4_26/TABLES/2

Yu, B., Chen, C., Zhou, H., Liu, B., & Ma, Q. (2020). GTB-PPI: Predict Protein–Protein Interactions Based on L1-Regularized Logistic Regression and Gradient Tree Boosting. Genomics, Proteomics & Bioinformatics, 18(5), 582–592. 10.1016/J.GPB.2021.01.001

Yu, W., Clyne, M., Khoury, M. J., & Gwinn, M. (2010). Phenopedia and Genopedia: disease-centered and gene-centered views of the evolving knowledge of human genetic associations. Bioinformatics, 26(1), 145–146. 10.1093/BIOINFORMATICS/BTP618

Zeng, X., Zhu, S., Liu, X., Zhou, Y., Nussinov, R., & Cheng, F. (2019). deepDR: a network-based deep learning approach to in silico drug repositioning. Bioinformatics, 35(24), 5191–5198. 10.1093/BIOINFORMATICS/BTZ418

Zhu, Y., Che, C., Jin, B., Zhang, N., Su, C., & Wang, F. (2020). Knowledge-driven drug repurposing using a comprehensive drug knowledge graph. Health Informatics Journal, 26(4), 2737–2750. 10.1177/1460458220937101

