## Supplement Document 1 for "An Integrated Knowledge Graph and Network Medicine Pipeline for Drug Repurposing: Benchmarking Across Human Diseases and Application to Amyotrophic Lateral Sclerosis"

#### ATCEA (ATC Category Enrichment Analysis) implementation

We performed ATCEA on drug candidates predicted by SAveRUNNER following these steps (Wray et al., 2018):

1. **Selection of Drugs:** We first selected all drugs that had associations with ALS ($p \neq 1$) in the SAveRUNNER result drug-disease network.
2. **ATC Classification:** These drugs were annotated using the Anatomical Therapeutic Chemical (ATC) classification system. We focused on the 3rd-level ATC codes (the first four characters), which represent therapeutic or pharmacological subgroups and provide a balanced level of specificity for enrichment analysis.
3. **Statistical Enrichment Test:** To evaluate whether certain ATC categories were overrepresented, we applied the Wilcoxon–Mann–Whitney test. This non-parametric statistical test compares the distribution of ranks of drugs within a specific ATC category to the distribution of ranks of drugs outside that category.
4. **Significance Assessment:** For each ATC category, we computed a p-value reflecting the probability of observing the given distribution by chance. Lower p-values suggest that drugs in the category tend to rank higher than expected.
5. **Multiple Testing Correction:** To correct for multiple hypothesis testing across ATC categories, we applied the Benjamini–Hochberg procedure (BH) to control the false discovery rate.
6. **Drug Prioritisation within Enriched Categories:** We selected drugs that met both of the following criteria: (i) statistical significance after Bonferroni correction in the SAveRUNNER analysis, and (ii) classification within ATC categories that were significantly enriched (BH-adjusted $p < 0.05$). These drugs were considered the promising repurposing candidates due to both strong network-based associations and pharmacological relevance.

#### Full ALS results from SAveRUNNER

**Table S1. Bonferroni-corrected significant ALS candidate drugs from SAveRUNNER across gene sets**

| Drug | p-value | Adjusted Similarity | Bonferroni-corrected p |
|---|---|---|---|
| **Known ALS Genes** | | | |
| (2e)-3-(3,4-dihydroxyphenyl)-2-iminopropanoic acid | 4.12623493619256e-06 | 0.9999541247739 | 0.0282853404876 |
| (5s)-5-iododihydro-2,4(1h,3h)-pyrimidinedione | 3.43605928894683e-06 | 0.9999541247739 | 0.02355418642573052 |
| 3-(4-hydroxyphenyl)-4,5-dihydro-5-isoxazole-acetic acid methyl ester | 1.87276694258551e-06 | 0.9999541247739 | 0.012837817391423671 |
| 3,4-dihydro-2h-pyrrolium-5-carboxylate | 7.06065591150591e-06 | 0.9999541247739 | 0.048400796273373015 |
| 4-hydroxyphenylpyruvic acid | 2.21750868385653e-06 | 0.9999541247739 | 0.015201022027836514 |
| 6s-5,6,7,8-tetrahydrobiopterin | 1.87276694258551e-06 | 0.9999541247739 | 0.012837817391423671 |
| aflibercept | 6.34391843948168e-07 | 0.999757584812291 | 0.004348756090264692 |
| amibegron | 5.99037140106353e-09 | 0.9999541247739 | 4.10639959542905e-05 |
| amobarbital | 2.06322153912111e-11 | 0.995906474185362 | 1.4143383650675208e-07 |
| aprobarbital | 1.25361174354056e-07 | 0.995906474185362 | 0.0008593508501970538 |
| barbital | 3.73868423252564e-10 | 0.995906474185362 | 2.562868041396326e-06 |
| bentazepam | 7.09024277860925e-06 | 0.955794953316456 | 0.04860361424736641 |
| butabarbital | 3.08479336347297e-16 | 0.988961789401233 | 2.114625850660721e-12 |
| butalbital | 4.330433910732e-09 | 0.987956930511467 | 2.968512445806786e-05 |

| | | | |
|---|---|---|---|
| butobarbital | 2.6440574615836703e-09 | 0.995906474185362 | 1.812501389915606e-05 |
| calcium levulinate | 2.26780024126438e-07 | 0.99944286995445 | 0.0015545770653867324 |
| cisatracurium | 6.0900790175243e-07 | 0.988961789401233 | 0.004174749166512908 |
| clomethiazole | 1.58716866629236e-06 | 0.9999541247739 | 0.010880041207434129 |
| copper | 5.26919305756484e-13 | 0.991905136253456 | 3.612031840960698e-09 |
| cytisine | 1.51287538216069e-08 | 0.99944286995445 | 0.0001037076074471153 |
| desflurane | 2.23243609875845e-06 | 0.96759319806178 | 0.015303349456989175 |
| dextromethorphan | 4.84444872482682e-06 | 0.993272288033523 | 0.03320869600868785 |
| dihydro-2-thioxo-5-((5-(2-(trifluoromethyl)phenyl)-2-furanyl)methyl)-4,6(1h,5h)-pyrimidinedione | 1.78171806792625e-10 | 0.995906474185362 | 1.2213677355634444e-06 |
| drotrecogin alfa | 2.23115093582435e-06 | 0.998488591365269 | 0.0152945396650759 2 |
| epibatidine | 3.22238243141478e-14 | 0.99944286995445 | 2.208943156734831e -10 |
| ethanol | 2.81882439983127e-10 | 0.95886443249193 | 1.9323041260843353e-06 |
| ethyl loflazepate | 2.89956175901036e-06 | 0.955794953316456 | 0.019876495858016017 |
| ferric maltol | 3.93421281733753e-06 | 0.9999541247739 | 0.026969028862848767 |
| ferrous succinate | 7.25939289227917e-06 | 0.995408666092403 | 0.04976313827657371 |
| flavin-adenine dinucleotide-n5-isobutyl ketone | 3.87291548348926e-06 | 0.9999541247739 | 0.026548835639318875 |
| flurazepam | 7.24716737324851e-07 | 0.955794953316456 | 0.004967933234361854 |
| gantacurium | 5.27242252521269e-07 | 0.9999541247739 | 0.003614245641033992 |
| glembatumumab vedotin | 2.3084757371916e-11 | 0.9999541247739 | 1.5824601178448418e-07 |
| glutathione | 1.75855831541008e-06 | 0.974539379415546 | 0.012054917252136099 |
| guvacine | 1.11961974201077e-08 | 0.993272288033523 | 7.674993331483829e-05 |
| heptabarbital | 3.9650072009473e-08 | 0.995906474185362 | 0.0002718012436249374 |
| hexobarbital | 2.93765765908709e-09 | 0.995906474185362 | 2.0137643253042005e-05 |
| imino-tryptophan | 5.99037140106353e-09 | 0.9999541247739 | 4.10639959542905e-05 |
| isoflurane | 1.74780610829304e-07 | 0.981939736118286 | 0.0011981210872348787 |
| lexacalcitol | 5.17727971880815e-06 | 0.9999541247739 | 0.035490252472429866 |
| lorazepam | 2.75901320130659e-06 | 0.955794953316456 | 0.018913035494956675 |
| medazepam | 4.94185773576123e-07 | 0.955794953316456 | 0.0033876434778643233 |
| metharbital | 9.35711816027234e-10 | 0.987956930511467 | 6.414304498866689e-06 |
| methoxyflurane | 1.80882047214498e-06 | 0.962017598897172 | 0.012399464336553838 |
| methylphenobarbital | 6.51777644997084e-09 | 0.995717129286109 | 4.467935756455011e-05 |
| n-isobutyl-n-[4-methoxyphenylsulfonyl]glycyl hydroxamic acid | 3.90279620432439e-06 | 0.998060651884434 | 0.026753667980643697 |
| nicotine | 2.77543415007286e-09 | 0.993272288033523 | 1.9025601098749458e-05 |
| pentobarbital | 1.46774653289431e-06 | 0.978033492099928 | 0.010061402482990495 |
| pentolinium | 6.195151814312721e-09 | 0.9999541247739 | 4.24677656871137e-05 |
| primidone | 1.41528961380685e-09 | 0.987956930511467 | 9.701810302645958e-06 |
| silver | 5.60338199462492e-06 | 0.985604558332078 | 0.038411183573153825 |
| sinapoyl coenzyme a | 6.63496460862948e-06 | 0.993272288033523 | 0.04548268239215508 |
| solabegron | 2.05229962402625e-07 | 0.9999541247739 | 0.0014068513922699945 |
| su-11652 | 9.918586161641189e-13 | 0.9999541247739 | 6.799190813805035e-09 |
| succinylcholine | 7.87440207775089e-07 | 0.993272288033523 | 0.005397902624298235 |
| talbutal | 6.95856863847653e-10 | 0.987956930511467 | 4.770098801675662e-06 |
| thiopental | 1.79773146010821e-06 | 0.989834507948104 | 0.01232344915904178 |

| | | | |
|---|---|---|---|
| thrombomodulin alfa | 1.82394880424258e-06 | 0.9999541247739 | 0.012503169053082887 |
| torcetrapib | 8.22166432190802e-07 | 0.9999541247739 | 0.005635950892667948 |
| varenicline | 4.3833084385905e-08 | 0.99944286995445 | 0.0003004757934653787 |
| vibegron | 2.05229962402625e-07 | 0.9999541247739 | 0.0014068513922699945 |
| voglibose | 1.64304476560519e-12 | 0.9999541247739 | 1.1263071868223577e-08 |
| zinc | 6.81002891827892e-07 | 0.98386883831092 | 0.0046682748234802 |
| zinc sulfate, unspecified form | 5.99458087181538e-06 | 0.984111038616434 | 0.04109285187629443 |
| **New Predicted ALS Genes** | | | |
| (1-benzyl-5-methoxy-2-methyl-1h-indol-3-yl)acetic acid | 7.66904909597165e-09 | 0.999954368904212 | 5.257133155288566e-05 |
| (2r,6s)-6-{[methyl(3,4,5-trimethoxyphenyl)amino]methyl}-1,2,5,6,7,8-hexahydroquinazoline-2,4-diamine | 5.17727971880815e-06 | 0.999954368904212 | 0.035490252472429866 |
| (2r)-({4-[amino(imino)methyl]phenyl}amino){5-ethoxy-2-fluoro-3-[(3r)-tetrahydrofuran-3-yloxy]phenyl}aceticacid | 3.41823552009093e-06 | 0.999954368904212 | 0.023432004490223327 |
| (2s,3r,4s,5s)-3,4-dihydroxy-2-[(methylsulfanyl)methyl]-5-(4-oxo-4,5-dihydro-1h-pyrrolo[3,2-d]pyrimidin-7-yl)pyrrolidinium | 7.22039941045998e-06 | 0.999954368904212 | 0.049495837958703166 |
| (3,4-dihydroxy-2-nitrophenyl)(phenyl)methanone | 5.45634574715296e-07 | 0.999954368904212 | 0.003740325009673354 |
| (3r)-3-hydroxydodecanoic acid | 3.41823552009093e-06 | 0.999954368904212 | 0.023432004490223327 |
| (4as,6r,8as)-11-[8-(1,3-dioxo-1,3-dihydro-2h-isoindol-2-yl)octyl]-6-hydroxy-3-methoxy-5,6,9,10-tetrahydro-4ah-[1]benzofuro[3a,3,2-ef][2]benzazepin-11-ium | 1.6982674071476e-07 | 0.999954368904212 | 0.0011641623075996797 |
| (9s)-9-[(8-ammoniooctyl)amino]-1,2,3,4,9,10-hexahydroacridinium | 3.26344004919557e-07 | 0.999954368904212 | 0.0022370881537235633 |
| (z)-2-[2-(4-methylpiperazin-1-yl)benzyl]diazenecarbothioamide | 7.66904909597165e-09 | 0.999954368904212 | 5.257133155288566e-05 |
| 1d09c3 | 4.43659025828176e-06 | 0.999344577843264 | 0.030412826220521463 |
| 2-[(5-hex-1-yn-1-ylfuran-2-yl)carbonyl]-n-methylhydrazinecarbothioamide | 5.45634574715296e-07 | 0.999954368904212 | 0.003740325009673354 |
| 2-[3-(5-mercapto-[1,3,4]thiadiazol-2yl)-ureido]-n-methyl-3-pentafluorophenyl-propionamide | 5.17727971880815e-06 | 0.999954368904212 | 0.035490252472429866 |
| 2-amino-3-(5-tert-butyl-3-(phosphonomethoxy)-4-isoxazolyl)propionic acid | 1.64304476560519e-12 | 0.999954368904212 | 1.1263071868223577e-08 |
| 3-(benzyloxy)pyridin-2-amine | 3.26344004919557e-07 | 0.998724639657417 | 0.0022370881537235633 |
| 3-indolebutyric acid | 3.92438826128784e-07 | 0.999954368904212 | 0.0026901681531128143 |
| 3,8-diamino-6-phenyl-5-[6-[1-[2-[(1,2,3,4-tetrahydro-9-acridinyl)amino]ethyl]-1h-1,2,3-triazol-5-yl]hexyl]-phenanthridinium | 3.26344004919557e-07 | 0.999954368904212 | 0.0022370881537235633 |
| 4-methoxyamphetamine | 5.3117548736126e-08 | 0.999693837916874 | 0.00036412079658614373 |
| 5-[2-(1h-pyrrol-1-yl)ethoxy]-1h-indole | 7.66904909597165e-09 | 0.999954368904212 | 5.257133155288566e-05 |
| 7,8-dihydroxy-4-phenyl-2h-chromen-2-one | 7.22039941045998e-06 | 0.999954368904212 | 0.049495837958703166 |
| abt-510 | 3.77488588904663e-08 | 0.999954368904212 | 0.000258768427694146 |
| alteplase | 2.67126749290815e-06 | 0.999758660113224 | 0.01831153866388537 |
| aminocaproic acid | 6.82382745143974e-06 | 0.999954368904212 | 0.04677733717961942 |
| amoxapine | 3.19018101950752e-08 | 0.984391632620964 | 0.0002186869088872404 |

| | | | |
|---|---|---|---|
| | | | 8 |
| amphetamine | 3.02291472299106e-13 | 0.999291168898622 | 2.0722080426103717e-09 |
| aniracetam | 4.93068143756522e-06 | 0.999758660113224 | 0.03379982125450959 |
| anti-inhibitor coagulant complex | 8.00922234366225e-08 | 0.999445093927621 | 0.0005490321916580473 |
| apd791 | 5.45634574715296e-07 | 0.999954368904212 | 0.003740325009673354 |
| arundic acid | 5.45634574715296e-07 | 0.999954368904212 | 0.003740325009673354 |
| benzbromarone | 1.14471640043162e-07 | 0.999954368904212 | 0.0007847030924958755 |
| benzoyl peroxide | 2.79684250600746e-06 | 0.99784370302018 | 0.01917235537868114 |
| bia | 6.63496460862948e-06 | 0.999954368904212 | 0.04548268239215508 |
| brasofensine | 4.79183276590321e-07 | 0.999954368904212 | 0.0032848013610266504 |
| brolucizumab | 2.08137521949324e-23 | 0.999954368904212 | 1.4267827129626163e-19 |
| butabarbital | 5.08406023190312e-07 | 0.981977549345431 | 0.003485123288969589 |
| butylscopolamine | 2.70997444742284e-06 | 0.999954368904212 | 0.01857687483708357 |
| captopril | 1.96313287848232e-13 | 0.999954368904212 | 1.3457275881996305e-09 |
| carglumic acid | 7.66904909597165e-09 | 0.999954368904212 | 5.257133155288566e-05 |
| carvedilol | 2.82318130405512e-06 | 0.998724639657417 | 0.01935290783929785 |
| chlorobutanol | 7.22039941045998e-06 | 0.999954368904212 | 0.049495837958703166 |
| chondroitin sulfate | 7.05670229602856e-09 | 0.999954368904212 | 4.837369423927578e-05 |
| coccidioides immitis spherule | 7.06065591150588e-06 | 0.999693837916874 | 0.048400796273372806 |
| copper | 1.32548214511933e-19 | 0.998102737873084 | 9.086180104793008e-16 |
| denibulin | 7.06065591150591e-06 | 0.999579561605748 | 0.048400796273373015 |
| dextromethorphan | 5.94420917025546e-08 | 0.997185090408625 | 0.0004074755386210118 |
| diacerein | 7.94075922614151e-12 | 0.999799431316062 | 5.4433904495200044e-08 |
| diethylene glycol diethyl ether | 5.17727971880815e-06 | 0.999954368904212 | 0.035490252472429866 |
| dihydrexidine | 1.6982674071476e-07 | 0.999954368904212 | 0.0011641623075996797 |
| droperidol | 6.63496460862948e-06 | 0.999954368904212 | 0.04548268239215508 |
| entacapone | 6.63496460862948e-06 | 0.999954368904212 | 0.04548268239215508 |
| ephedra sinica root | 6.09033904435271e-07 | 0.998724639657417 | 0.004174927414903783 |
| eprosartan | 7.22039941045998e-06 | 0.999954368904212 | 0.049495837958703166 |
| estradiol | 6.88024084269195e-08 | 0.999827031045199 | 0.0004716405097665332 |
| ethinylestradiol | 7.22039941045998e-06 | 0.999954368904212 | 0.049495837958703166 |
| etoperidone | 6.78223711647784e-06 | 0.994703565892634 | 0.04649223543345559 |
| fexaramine | 4.79183276590321e-07 | 0.999954368904212 | 0.0032848013610266504 |
| flavin adenine dinucleotide | 1.14234062627064e-06 | 0.99409232180717 | 0.007830744993085238 |
| foreskin keratinocyte (neonatal) | 6.36114691115641e-06 | 0.999267651323477 | 0.04360566207597719 |
| gamma-glutamylcysteine | 1.64304476560519e-12 | 0.999954368904212 | 1.1263071868223577e-08 |
| ginseng | 4.33068693757282e-06 | 0.999954368904212 | 0.02968685895706168 |
| glutathione | 6.50095213951826e-15 | 0.996573444072696 | 4.4564026916397674e-11 |
| halofantrine | 5.45634574715296e-07 | 0.999954368904212 | 0.003740325009673354 |
| haloperidol | 5.92824876073198e-07 | 0.996138030784651 | 0.0040638145254817725 |
| hyaluronidase (human recombinant) | 7.22039941045998e-06 | 0.999954368904212 | 0.049495837958703166 |
| hyaluronidase (ovine) | 7.66904909597165e-09 | 0.999954368904212 | 5.257133155288566e-05 |
| hydroxy(oxo)(2-{(1s)-2,2,2-trifluoro-1-[2 (trimethylarsonio)ethoxy]ethyl}phenyl)ammonium | 3.41823552009093e-06 | 0.999954368904212 | 0.023432004490223327 |
| irbesartan | 7.22039941045998e-06 | 0.999954368904212 | 0.049495837958703166 |
| isoniazid | 5.41879216071394e-08 | 0.999954368904212 | 0.0003714582026169406 |
| itopride | 1.57643723402687e-09 | 0.999954368904212 | 1.0806477239254195e-05 |
| levacetylmethadol | 6.06345181456911e-06 | 0.999954368904212 | 0.04156496218887125 |
| licofelone | 6.25579659862704e-07 | 0.999954368904212 | 0.004288348568358836 |
| loratadine | 2.76396560310874e-06 | 0.999954368904212 | 0.018946984209310413 |
| maprotiline | 5.69090212139854e-06 | 0.994990537909104 | 0.03901113404218699 |
| mesoridazine | 5.45634574715296e-07 | 0.999954368904212 | 0.003740325009673354 |
| methotrimeprazine | 5.53752271508017e-07 | 0.995661201142903 | 0.0037959718211874566 |
| mmda | 7.52004243004547e-07 | 0.999693837916874 | 0.005154989085796169 |
| myrrh | 1.21327339352332e-08 | 0.999954368904212 | 8.316989112602359e-05 |
| n-dodecyl-n,n-dimethyl-3-ammonio-1- | 2.67126749290815e-06 | 0.999954368904212 | 0.01831153866388537 |

| | | | |
|---|---|---|---|
| propanesulfonate | | | |
| n(5)-[(hydroxyamino)(imino)methyl]-l-ornithine | 6.75902614203278e-06 | 0.999954368904212 | 0.04633312420363471 |
| nadph | 3.81160565925868e-06 | 0.999954368904212 | 0.02612855679421825 |
| nap-226-90 | 3.26344004919557e-07 | 0.999954368904212 | 0.0022370881537235633 |
| neovastat | 1.69050822537905e-06 | 0.999954368904212 | 0.011588433884973388 |
| nimesulide | 2.05229962402625e-07 | 0.999954368904212 | 0.0014068513922699945 |
| nomifensine | 4.21592227846596e-08 | 0.999693837916874 | 0.00028900147218884I5 |
| norepinephrine | 2.06978703549163e-07 | 0.999057588192694 | 0.0014188390128295125 |
| nortriptyline | 2.31577344684966e-08 | 0.995319090397961 | 0.0001587462697815442 |
| ocaperidone | 3.12685467769477e-06 | 0.998274265555095 | 0.02143458881559765 |
| olanzapine | 5.5961741157027e-08 | 0.990255756305885 | 0.0003836177356314201 |
| oleandrin | 1.54286877355452e-09 | 0.999911156958969 | 1.0576365442716236e-05 |
| omega-3-acid ethyl esters | 2.08137521949324e-23 | 0.999954368904212 | 1.4267827129626163e-19 |
| paclitaxel | 1.31559608341128e-06 | 0.999861476742928 | 0.009018411151784325 |
| pentobarbital | 1.59830239184147e-06 | 0.984190605225789 | 0.010956362896073277 |
| perospirone | 1.45163499248401e-06 | 0.999445093927621 | 0.009950957873477889 |
| pipamperone | 2.92941300036266e-08 | 0.999344577843264 | 0.00020081126117486034 |
| porphobilinogen | 6.63496460862948e-06 | 0.999954368904212 | 0.04548268239215508 |
| potassium nitrate | 2.08137521949324e-23 | 0.999954368904212 | 1.4267827129626163e-19 |
| prasterone | 4.0858514777213e-06 | 0.989703082686074 | 0.028008511879779513 |
| propiomazine | 2.20667822850264e-06 | 0.99751978147876 | 0.015126779256385598 |
| px-12 | 5.45634574715296e-07 | 0.999954368904212 | 0.003740325009673354 |
| quisqualic acid | 3.43605928894683e-06 | 0.999954368904212 | 0.02355418642573052 |
| ranibizumab | 1.64304476560519e-12 | 0.999954368904212 | 1.1263071868223577e-08 |
| remoxipride | 6.11244904471688e-07 | 0.999827031045199 | 0.004190083820153422 |
| resveratrol | 1.54019254515067e-09 | 0.999445093927621 | 1.0558019897007844e-05 |
| rilonacept | 8.98211291370462e-07 | 0.999954368904212 | 0.006157238402344517 |
| rilpivirine | 7.66904909597165e-09 | 0.999954368904212 | 5.257133155288566e-05 |
| ropinirole | 1.57922538049435e-06 | 0.998154337776626 | 0.01082558998328877 |
| sertindole | 4.29297896488751e-08 | 0.999445093927621 | 0.0002942837080430388 |
| succinylcholine | 1.28914650697306e-06 | 0.99751978147876 | 0.008837099305300326 |
| sulpiride | 1.53605624219329e-06 | 0.999344577843264 | 0.010529665540235003 |
| tacrine(8)-4-aminoquinoline | 9.40860846467867e-10 | 0.999954368904212 | 6.449601102537229e-06 |
| talbutal | 4.85766058343711e-06 | 0.983944799433257 | 0.03329926329946139 |
| taurocholic acid | 5.45634574715296e-07 | 0.999954368904212 | 0.003740325009673354 |
| tecastemizole | 7.22039941045998e-06 | 0.999954368904212 | 0.049495837958703166 |
| terazosin | 5.42601097651533e-07 | 0.999579561605748 | 0.003719530524401259 |
| tetrahydropalmatine | 5.98117327209416e-06 | 0.999954368904212 | 0.04100094278020546 |
| thiethylperazine | 6.63496460862948e-06 | 0.999954368904212 | 0.04548268239215508 |
| thioproperazine | 2.01833844288692e-09 | 0.999954368904212 | 1.3835710025989838e-05 |
| thioridazine | 2.09825617828286e-15 | 0.999954368904212 | 1.4383546102129008e-11 |
| thiothixene | 3.49352810007844e-08 | 0.999954368904212 | 0.000239481351260377 |
| thrombin alfa | 2.5947360855424e-07 | 0.999344577843264 | 0.0017786915866393152 |
| trimipramine | 6.19156608875264e-06 | 0.993290093599181 | 0.04244318553839935 |
| valsartan | 5.45634574715296e-07 | 0.999954368904212 | 0.003740325009673354 |
| vanoxerine | 1.14660213573041e-06 | 0.999579561605748 | 0.00785995764043196 |
| willardiine | 7.22039941045998e-06 | 0.999954368904212 | 0.049495837958703166 |
| ykp-1358 | 5.45634574715296e-07 | 0.999954368904212 | 0.003740325009673354 |
| zinc | 5.19944827556156e-21 | 0.996212914302392 | 3.56422179289745e-17 |
| zinc acetate | 3.43528126038286e-17 | 0.996212914302392 | 2.3548853039924504e-13 |
| zinc chloride | 5.59017237716667e-14 | 0.996963869642081 | 3.8320631645477525e-10 |
| zinc sulfate, unspecified form | 4.75973664739322e-16 | 0.996963869642081 | 3.262799471788052e-12 |
| **All ALS Genes** | | | |
| (2s)-2-ammonio-3-[5-(2-methyl-2-propanyl)-3-oxido-1,2-oxazol-4-yl]propanoate | 7.22039941045998e-06 | 0.999954368231161 | 0.049495837958703166 |
| (4-fluorophenyl)(pyridin-4-yl)methanone | 3.41823552009093e-06 | 0.999954368231161 | 0.023432004490223327 |

| | | | |
|---|---|---|---|
| [2'-hydroxy-3'-(1h-pyrrolo[3,2-c]pyridin-2-yl)-biphenyl-3-ylmethyl]-urea | 1.27821325084469e-06 | 0.999954368231161 | 0.00876215183454035 |
| 1-[3-({[(4-amino-5-fluoro-2-methylquinolin-3-yl)methyl]thio}methyl)phenyl]-2,2,2-trifluoroethane-1,1-diol | 9.918586161641189e-13 | 0.999954368231161 | 6.799190813805035e-09 |
| 1-azepan-1-yl-2-phenyl-2-(4-thioxo-1,4-dihydro-pyrazolo[3,4-d]pyrimidin-5-yl)ethanone adduct | 7.22039941045998e-06 | 0.999954368231161 | 0.0494958379587031666 |
| 12-phenylheme | 5.45634574715296e-07 | 0.999954368231161 | 0.003740325009673354 |
| 2-[(5-hex-1-yn-1-ylfuran-2-yl)carbonyl]-n-methylhydrazinecarbothioamide | 7.22039941045998e-06 | 0.999954368231161 | 0.0494958379587031666 |
| 2-amino-3-(5-tert-butyl-3-(phosphonomethoxy)-4-isoxazolyl)propionic acid | 7.22039941045998e-06 | 0.999954368231161 | 0.0494958379587031666 |
| 3-[(4'-cyanobiphenyl-4-yl)oxy]-n-hydroxypropanamide | 6.63496460862948e-06 | 0.999954368231161 | 0.04548268239215508 |
| 3-indolebutyric acid | 6.25579659862704e-07 | 0.999954368231161 | 0.004288348568358836 |
| 3-tyrosine | 6.63496460862948e-06 | 0.999954368231161 | 0.04548268239215508 |
| 3,6,9,12,15-pentaoxaheptadecane | 6.11031905461104e-09 | 0.999954368231161 | 4.188623711935868e-05 |
| 4-amino-n-[4-(benzyloxy)phenyl]butanamide | 3.09203286925253e-09 | 0.999954368231161 | 2.1195885318726095e-05 |
| 4-methoxyamphetamine | 6.281291516160711e-09 | 0.999693834262151 | 4.305825334328167e-05 |
| 4,7-dioxosebacic acid | 4.79183276590321e-07 | 0.999954368231161 | 0.0032848013610266504 |
| 5-(hexahydro-2-oxo-1h-thieno[3,4-d]imidazol-6-yl)pentanal | 1.64304476560519e-12 | 0.999954368231161 | 1.1263071868223577e-08 |
| 7,8-dihydroxy-4-phenyl-2h-chromen-2-one | 7.22039941045998e-06 | 0.999954368231161 | 0.0494958379587031666 |
| 9-deazainosine-2',3'-o-ethylidenephosphonate | 7.66904909597165e-09 | 0.999954368231161 | 5.257133155288566e-05 |
| abt-510 | 2.66130888158427e-13 | 0.999954368231161 | 1.824327238326017e-09 |
| acepromazine | 1.31559608341128e-06 | 0.999579556784254 | 0.009018411151784325 |
| alfimeprase | 2.85977424710478e-06 | 0.999954368231161 | 0.01960375246390327 |
| amoxapine | 5.1353314596074e-07 | 0.984391538649588 | 0.0035202697155608726 |
| amphetamine | 5.50661389802245e-13 | 0.999291161318564 | 3.774783827094389e-09 |
| angiotensin ii | 5.45634574715296e-07 | 0.999954368231161 | 0.003740325009673354 |
| aniracetam | 2.33120800222796e-08 | 0.999758657147371 | 0.000159804308552726 |
| anti-inhibitor coagulant complex | 3.26344004919557e-07 | 0.999445087792178 | 0.0022370881537235633 |
| arundic acid | 7.22039941045998e-06 | 0.999954368231161 | 0.0494958379587031666 |
| benzoyl peroxide | 3.08164319309904e-07 | 0.998480558845047 | 0.0021124664088693922 |
| beta-d-fructofuranose 1,6-bisphosphate | 7.22039941045998e-06 | 0.999954368231161 | 0.0494958379587031666 |
| beta-l-fucose | 3.26344004919557e-07 | 0.999954368231161 | 0.0022370881537235633 |
| bevasiranib | 5.45634574715296e-07 | 0.999954368231161 | 0.003740325009673354 |
| bl-1020 | 2.31208795561913e-07 | 0.999954368231161 | 0.0015849362935769135 |
| brolucizumab | 5.45634574715296e-07 | 0.999954368231161 | 0.003740325009673354 |
| butabarbital | 5.71610450305168e-07 | 0.981977444926205 | 0.003918389636841927 |
| butylscopolamine | 1.61895395041205e-08 | 0.999954368231161 | 0.000110979293300746 |
| canaline | 1.64304476560519e-12 | 0.999954368231161 | 1.1263071868223577e-08 |
| cannabidiol | 1.52822358612671e-06 | 0.985019054016815 | 0.010475972682898597 |
| captopril | 8.24824204944544e-09 | 0.999954368231161 | 5.65416992489485e-05 |
| carboxymethylcellulose | 7.66904909597165e-09 | 0.999954368231161 | 5.257133155288566e-05 |
| carglumic acid | 1.64304476560519e-12 | 0.999954368231161 | 1.1263071868223577e-08 |
| carvedilol | 3.72655744213597e-08 | 0.998724627131966 | 0.0002554555126584207 |
| cerliponase alfa | 7.22039941045998e-06 | 0.999954368231161 | 0.0494958379587031666 |
| chlorobutanol | 7.22039941045998e-06 | 0.999954368231161 | 0.0494958379587031666 |
| chondroitin sulfate | 2.95998943486044e-12 | 0.999954368231161 | 2.0290727575968314e-08 |
| coccidioides immitis spherule | 3.81624385081978e-08 | 0.999693834262151 | 0.0002616035159736959 |
| copper | 3.5675660213506e-24 | 0.998194590465183 | 2.4455665076358363e-20 |

| | | | |
|---|---|---|---|
| desmethylsertraline | 7.22039941045998e-06 | 0.999954368231161 | 0.049495837958703166 |
| dexniguldipine | 5.17727971880815e-06 | 0.999954368231161 | 0.035490252472429866 |
| dextromethorphan | 9.77686557329946e-07 | 0.997185066091787 | 0.00670204135049678 |
| diacerein | 2.33081699577002e-09 | 0.999799428796374 | 1.597775050600349e-05 |
| dimetindene | 9.01450609253643e-08 | 0.999954368231161 | 0.0006179443926433723 |
| docetaxel | 5.09411431011238e-06 | 0.999861474926924 | 0.03492015359582036 |
| doxazosin | 6.31617032386219e-06 | 0.999344570758196 | 0.04329734757007531 |
| droperidol | 5.00184201573872e-06 | 0.999954368231161 | 0.0342876270178893 |
| droxidopa | 5.32655939377505e-06 | 0.998724627131966 | 0.036513564644327964 |
| ephedra sinica root | 1.91021926296575e-06 | 0.998724627131966 | 0.013094553047630216 |
| eprosartan | 7.66904909597165e-09 | 0.999954368231161 | 5.257133155288566e-05 |
| estradiol | 2.05345979923519e-06 | 0.999827028834387 | 0.014076466923757228 |
| etiprednol dicloacetate | 5.17727971880815e-06 | 0.999954368231161 | 0.035490252472429866 |
| fexaramine | 6.63496460862948e-06 | 0.999954368231161 | 0.0454826823921550 8 |
| fg-9041 | 7.66904909597165e-09 | 0.999954368231161 | 5.257133155288566e-05 |
| flavin adenine dinucleotide | 1.06487464645732e-06 | 0.995421742786652 | 0.007299715701464929 |
| gentamicin | 6.11031905461104e-09 | 0.999954368231161 | 4.188623711935868e-05 |
| glutathione | 9.30173227616133e-14 | 0.997136509639809 | 6.376337475308592e-10 |
| glycolic acid | 7.22039941045998e-06 | 0.999954368231161 | 0.049495837958703166 |
| hydroxyzine | 2.59474300611377e-06 | 0.999954368231161 | 0.01778696330690989 3 |
| inosine | 5.45634574715296e-07 | 0.999954368231161 | 0.003740325009673354 |
| iodo-willardiine | 7.22039941045998e-06 | 0.999954368231161 | 0.049495837958703166 |
| irbesartan | 6.63496460862948e-06 | 0.999954368231161 | 0.0454826823921550 8 |
| isoniazid | 2.40043272097688e-10 | 0.999954368231161 | 1.6454966302296514e-06 |
| itopride | 1.94500523834455e-06 | 0.999954368231161 | 0.013333010908851889 |
| kelatorphan | 6.63496460862948e-06 | 0.999954368231161 | 0.0454826823921550 8 |
| laevulinic acid | 6.63496460862948e-06 | 0.999954368231161 | 0.0454826823921550 8 |
| losartan | 7.66904909597165e-09 | 0.999954368231161 | 5.257133155288566e-05 |
| melperone | 6.63496460862948e-06 | 0.999954368231161 | 0.0454826823921550 8 |
| mesoridazine | 2.16610738409332e-08 | 0.999954368231161 | 0.000148486661179597 |
| methotrimeprazine | 7.09270639991479e-08 | 0.995661166487296 | 0.000486205023714158 |
| methylphosphinate | 6.63496460862948e-06 | 0.999954368231161 | 0.0454826823921550 8 |
| methylthioninium | 5.17727971880815e-06 | 0.999954368231161 | 0.035490252472429866 |
| metreleptin | 7.66904909597165e-09 | 0.999954368231161 | 5.257133155288566e-05 |
| mf268 | 5.17727971880815e-06 | 0.999954368231161 | 0.035490252472429866 |
| miltefosine | 6.63496460862948e-06 | 0.999954368231161 | 0.0454826823921550 8 |
| mirtazapine | 1.72939114773657e-06 | 0.998724627131966 | 0.011854976317734188 |
| mln-977 | 7.22039941045998e-06 | 0.999954368231161 | 0.049495837958703166 |
| mmda | 1.50102371949989e-08 | 0.999693834262151 | 0.000102895175971717 |
| myrrh | 2.71161453977822e-07 | 0.999954368231161 | 0.001858811767017 97 |
| n-[1-(4-carbamimidoyl-benzylcarbamoyl)-3-methylsulfanyl-propyl]-3-hydroxy-2-propoxyamino-butyramid | 3.26344004919557e-07 | 0.999954368231161 | 0.0022370881537235633 |
| n-acetyl-alpha-d-glucosamine | 3.27819135554802e-06 | 0.999758657147371 | 0.02247200174228168 |
| n(5)-[(hydroxyamino)(imino)methyl]-l-ornithine | 5.55877005308628e-06 | 0.999954368231161 | 0.03810536871390645 |
| nadph | 2.36834841178667e-06 | 0.999954368231161 | 0.016235028362797624 |
| neovastat | 3.47677468787665e-07 | 0.999954368231161 | 0.0023833290485394437 |
| nialamide | 1.16138060962492e-06 | 0.999954368231161 | 0.007961264078978826 |
| nomifensine | 3.39657328642927e-07 | 0.999693834262151 | 0.002328350987847265 |
| nortriptyline | 4.83209262781869e-07 | 0.995319053546835 | 0.003312399496369712 |
| ocaperidone | 6.407537460919e-06 | 0.998274249384162 | 0.043923669294599746 |
| odevixibat | 7.22039941045998e-06 | 0.999954368231161 | 0.049495837958703166 |
| olanzapine | 5.85753050085254e-08 | 0.99025569049673 | 0.0004015337158334416 |
| oleandrin | 8.89671139787346e-10 | 0.999911155735963 | 6.098695663242257e-06 |
| paclitaxel | 4.61414161566602e-06 | 0.999861474926924 | 0.031629940775390566 |
| perospirone | 1.29650648587913e-07 | 0.999445087792178 | 0.0008887551960701436 |

| | | | |
|---|---|---|---|
| pipamperone | 2.09887088796777e-07 | 0.999344570758196 | 0.0014387759937019063 |
| ranibizumab | 6.63496460862948e-06 | 0.999954368231161 | 0.04548268239215508 |
| remoxipride | 4.12623493619254e-06 | 0.999827028834387 | 0.028285340487599862 |
| resveratrol | 1.18519680531635e-08 | 0.999445087792178 | 8.12452410044358e-05 |
| rilonacept | 4.72759838003071e-06 | 0.999954368231161 | 0.03240768689511052 |
| saprisartan | 1.64304476560519e-12 | 0.999954368231161 | 1.1263071868223577e-08 |
| sertindole | 6.55686965686697e-07 | 0.999445087792178 | 0.0044947341497823074 |
| silodosin | 4.95163289958701e-09 | 0.999344570758196 | 3.3943443526668954e-05 |
| sri-9439 | 1.6982674071476e-07 | 0.999954368231161 | 0.0011641623075996797 |
| succinylcholine | 4.71430135002507e-06 | 0.997519759582574 | 0.03231653575442186 |
| sucralfate | 7.14083943115089e-10 | 0.999827028834387 | 4.895045430053935e-06 |
| sulpiride | 1.22752442698352e-06 | 0.999344570758196 | 0.00841467994697203 |
| talbutal | 1.19833171621499e-06 | 0.983944703484201 | 0.008214563914653757 |
| tariquidar | 3.41823552009093e-06 | 0.999954368231161 | 0.023432004490223327 |
| tecastemizole | 7.66904909597165e-09 | 0.999954368231161 | 5.257133155288566e-05 |
| terazosin | 8.82356916610134e-08 | 0.999579556784254 | 0.0006048556663362469 |
| tetrahydropalmatine | 7.78882241632094e-07 | 0.999954368231161 | 0.005339237766388004 |
| thioproperazine | 7.95518978560357e-15 | 0.999954368231161 | 5.4532825980312475e-11 |
| thioridazine | 7.35603329534109e-13 | 0.999954368231161 | 5.042560823956317e-09 |
| thiothixene | 8.9135057394078e-10 | 0.999954368231161 | 6.110208184364047e-06 |
| thrombomodulin alfa | 4.12623493619256e-06 | 0.999954368231161 | 0.0282853404876 |
| tropifexor | 1.64304476560519e-12 | 0.999954368231161 | 1.1263071868223577e-08 |
| vanoxerine | 1.16347421886965e-06 | 0.999579556784254 | 0.007975615770351451 |
| veglin | 5.45634574715296e-07 | 0.999954368231161 | 0.003740325009673354 |
| viloxazine | 2.55975201975125e-06 | 0.997252753370356 | 0.01754710009539482 |
| volixibat | 5.8282486438352e-06 | 0.999954368231161 | 0.039952644453490296 |
| willardiine | 7.22039941045998e-06 | 0.999954368231161 | 0.049495837958703166 |
| ykp-1358 | 1.6982674071476e-07 | 0.999954368231161 | 0.0011641623075996797 |
| zidovudine diphosphate | 7.22039941045998e-06 | 0.999954368231161 | 0.049495837958703166 |
| zinc | 1.91331490001255e-14 | 0.996212883276447 | 1.311577363958603e-10 |
| zinc acetate | 1.16446091072917e-15 | 0.996212883276447 | 7.982379543048461e-12 |
| zinc chloride | 1.55896026249452e-18 | 0.996963843758973 | 1.0686672599399933e-14 |
| zinc sulfate, unspecified form | 4.59469191953336e-15 | 0.996963843758973 | 3.149661310840119e-11 |

### Reference

Wray, N. R., Ripke, S., Mattheisen, M., Trzaskowski, M., Byrne, E. M., Abdellaoui, A., … Sullivan, P. F. (2018). Genome-wide association analyses identify 44 risk variants and refine the genetic architecture of major depression. *Nature Genetics 2018 50:5*, *50*(5), 668–681. doi: 10.1038/s41588-018-0090-3
