## Supplement Document 5 for "An Integrated Knowledge Graph and Network Medicine Pipeline for Drug Repurposing: Benchmarking Across Human Diseases and Application to Amyotrophic Lateral Sclerosis"

### als drug repurposing ehr evaluation

generated: 2026-03-27 11:45

#### study summary

- study: als drug repurposing ehr evaluation
- disease: als
- diagnosis event: Neurology Motor Nerve Clinic Letter
- candidate drugs: 15
- matching ratio: 5:1
- age caliper: +/- 5 years
- fdr correction: fdr_bh (alpha=0.05)

##### cohort

- total patients: 2,200
- deceased: 1,495
- alive (censored): 705

#### drug exposure

15 of 15 candidate drugs found in ehr records.

| drug | n_patients | n_prescriptions |
| --- | --- | --- |
| lorazepam | 62 | 185 |
| haloperidol | 39 | 72 |
| losartan | 29 | 119 |
| doxazosin | 27 | 150 |
| mirtazapine | 24 | 110 |
| loratadine | 15 | 55 |
| irbesartan | 9 | 19 |
| olanzapine | 5 | 17 |
| ropinirole | 3 | 18 |
| valsartan | 3 | 11 |
| carvedilol | 2 | 4 |
| nortriptyline | 2 | 14 |
| captopril | 1 | 3 |
| norepinephrine | 1 | 1 |
| terazosin | 1 | 7 |

##### timing

14 drugs with pre-diagnosis use.

| drug | n_patients_total | n_pre_diagnosis | n_post_diagnosis | n_pre_only | n_post_only | n_both | n_no_diagnosis |
| --- | --- | --- | --- | --- | --- | --- | --- |
| losartan | 29 | 19 | 14 | 15 | 10 | 4 | 0 |

| drug | n_patients_total | n_pre_diagnosis | n_post_diagnosis | n_pre_only | n_post_only | n_both | n_no_diagnosis |
|---|---|---|---|---|---|---|---|
| doxazosin | 27 | 15 | 17 | 10 | 12 | 5 | 0 |
| lorazepam | 62 | 9 | 55 | 7 | 53 | 2 | 0 |
| loratadine | 15 | 6 | 12 | 3 | 9 | 3 | 0 |
| irbesartan | 9 | 6 | 3 | 6 | 3 | 0 | 0 |
| mirtazapine | 24 | 5 | 20 | 4 | 19 | 1 | 0 |
| haloperidol | 39 | 4 | 36 | 3 | 35 | 1 | 0 |
| ropinirole | 3 | 2 | 2 | 1 | 1 | 1 | 0 |
| valsartan | 3 | 1 | 3 | 0 | 2 | 1 | 0 |
| norepinephrine | 1 | 1 | 0 | 1 | 0 | 0 | 0 |
| captopril | 1 | 1 | 0 | 1 | 0 | 0 | 0 |
| carvedilol | 2 | 1 | 1 | 1 | 1 | 0 | 0 |
| nortriptyline | 2 | 1 | 1 | 1 | 1 | 0 | 0 |
| olanzapine | 5 | 1 | 4 | 1 | 4 | 0 | 0 |
| terazosin | 1 | 0 | 1 | 0 | 1 | 0 | 0 |

#### pre-diagnosis survival analysis

**what is this analysis?**

for each candidate drug, we find patients who were prescribed it *before* their first diagnosis event (e.g. first neurology motor nerve clinic letter). these are patients who happened to already be taking the drug for another condition before they developed als/mnd.

we then **match** each drug user to similar patients who *never* took that drug, using propensity-score-style matching on:

- **age at diagnosis** (within the caliper — e.g. ±2 years)
- **sex** (exact match)
- **concurrent medication count** (proxy for overall health burden)
- **calendar year of diagnosis** (controls for changes in care over time)

**interpreting the hazard ratio (hr):**

- **hr < 1.0** → drug users survived *longer* than matched controls (potentially protective)
- **hr = 1.0** → no difference in survival
- **hr > 1.0** → drug users survived *shorter* than matched controls (potentially harmful, or confounding by indication)

**multiple testing correction:** when testing many drugs simultaneously, some will appear significant by chance alone. we apply false discovery rate (fdr) correction to account for this. the "significant_fdr" column indicates whether the result survives this correction.

**important caveat:** this is an observational study, not a randomised trial. a significant association does not prove causation — the drug might be a marker for an underlying condition that independently affects survival.

analysed 7 drugs with sufficient data.

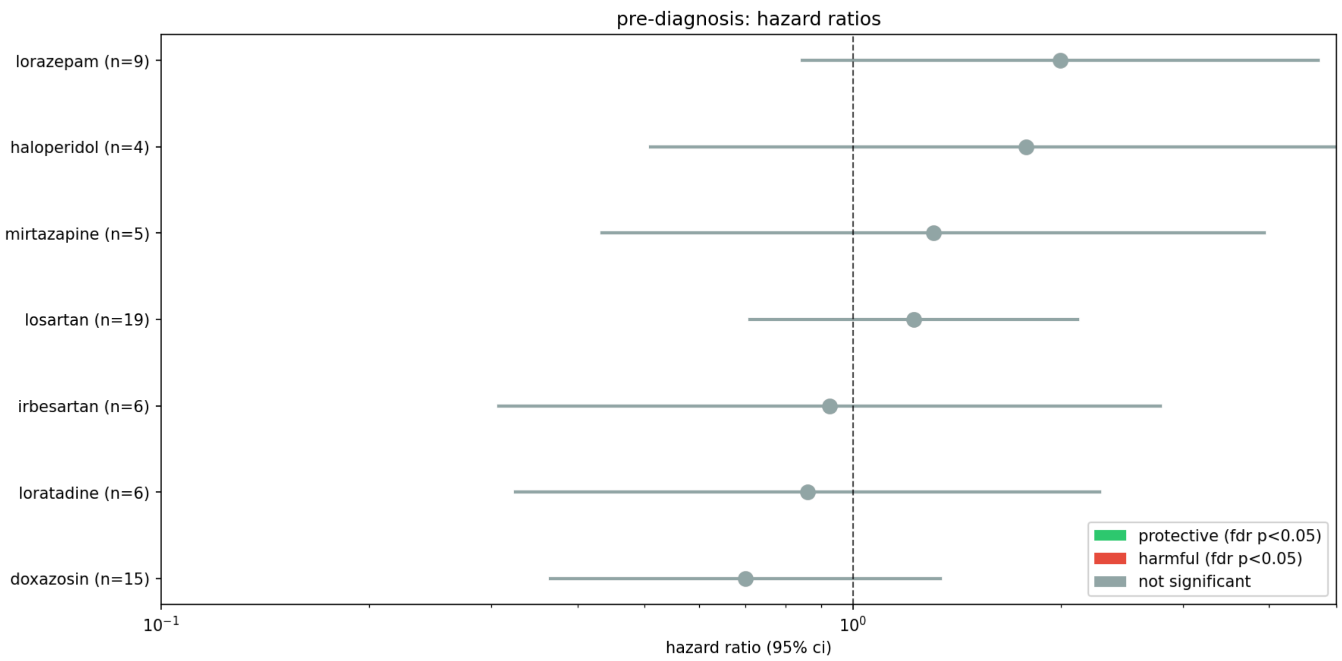


| drug | n_users | n_controls | n_users_deceased | n_controls_deceased | median_survival_users | median_survival_controls | hazard_ratio | hr_ci_lo |
|---|---|---|---|---|---|---|---|---|
| doxazosin | 15 | 75 | 11 | 60 | 1.539 | 1.005 | 0.699 | 0.365 |
| loratadine | 6 | 30 | 5 | 26 | 0.756 | 0.862 | 0.860 | 0.325 |
| irbesartan | 6 | 30 | 4 | 19 | 12.339 | 3.510 | 0.925 | 0.307 |
| losartan | 19 | 95 | 16 | 71 | 1.309 | 1.656 | 1.226 | 0.711 |
| mirtazapine | 5 | 25 | 4 | 20 | 1.758 | 1.243 | 1.308 | 0.434 |
| haloperidol | 4 | 20 | 3 | 16 | 0.162 | 0.868 | 1.781 | 0.510 |
| lorazepam | 9 | 45 | 7 | 28 | 1.780 | 2.448 | 1.996 | 0.845 |

#### kaplan-meier survival curves — all drugs with viable results

these plots show the estimated survival probability over time for drug users (red) vs matched controls (blue). the shaded bands are 95% confidence intervals. if the red curve stays above the blue curve, it means drug users survived longer. the text box in each plot shows the hazard ratio, p-value, and sample sizes.

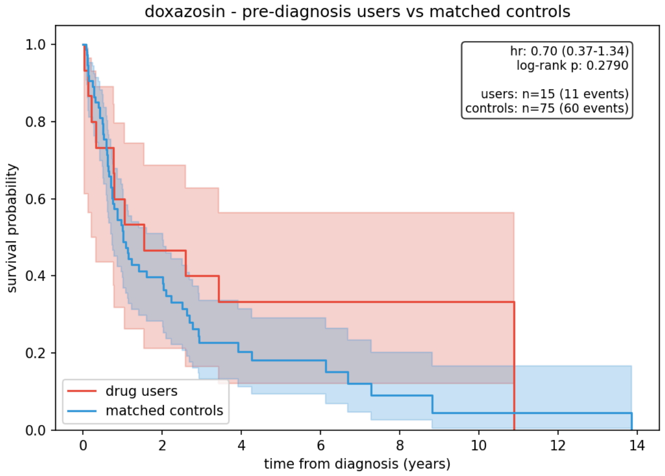


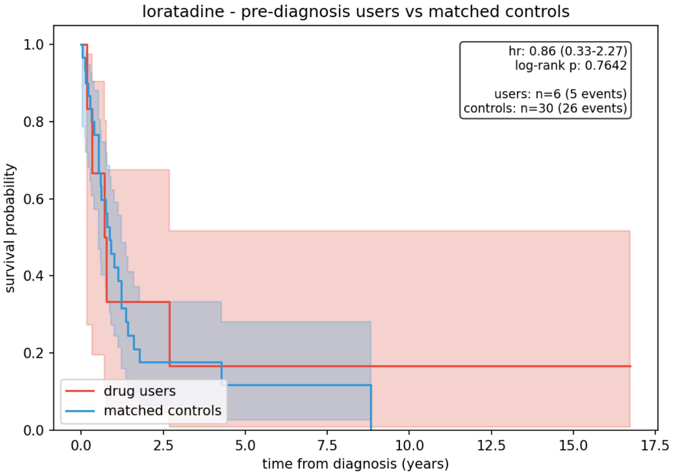


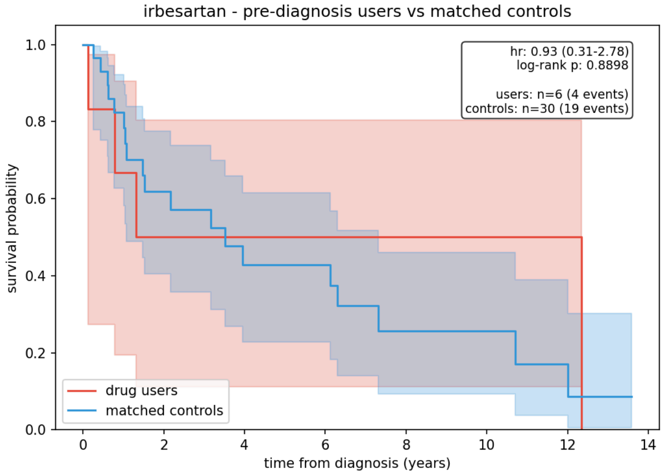


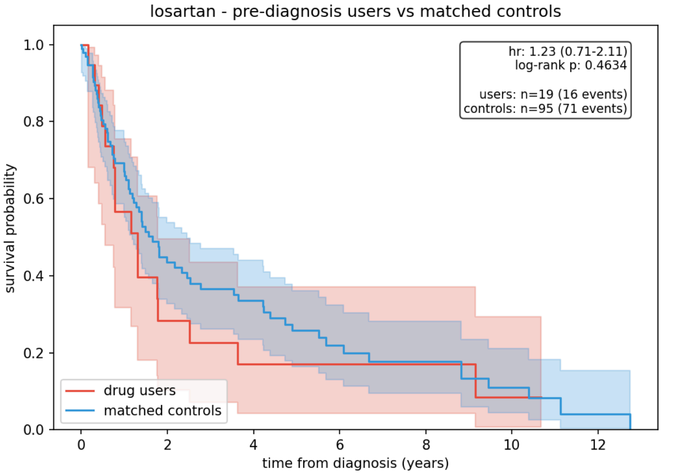


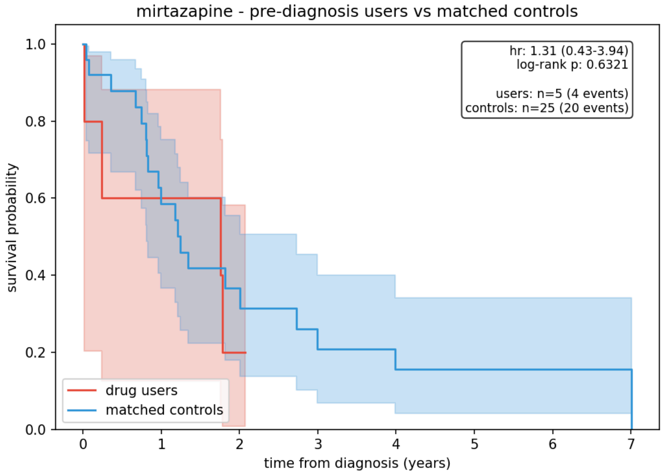


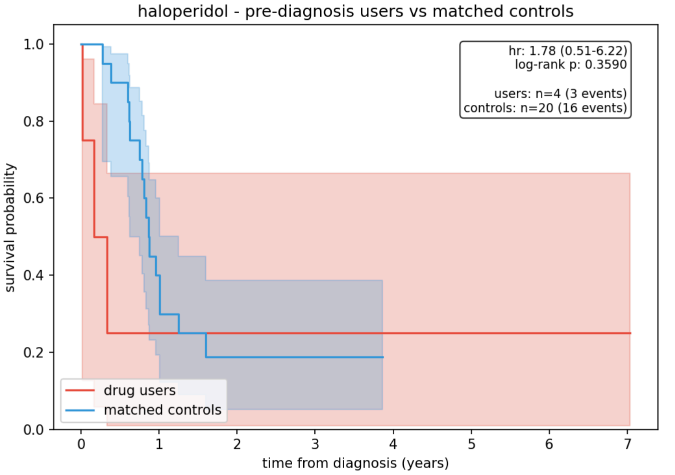


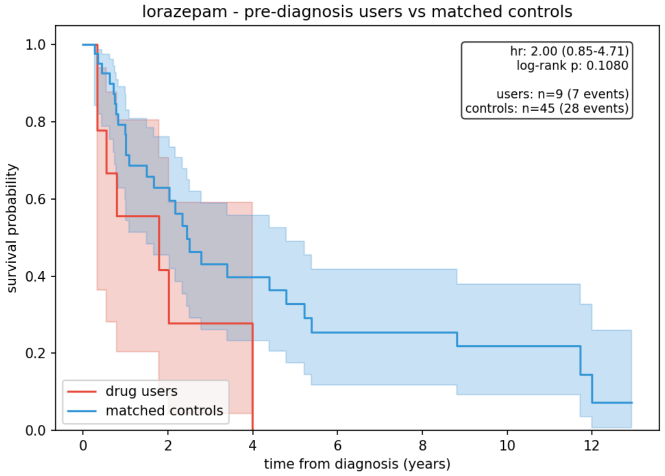


#### post-diagnosis survival analysis (landmark)

**what is landmark analysis?**

when we look at drugs prescribed *after* diagnosis, there is a fundamental bias called **immortal time bias**: patients who died shortly after diagnosis never had the chance to receive a new prescription, so they automatically end up in the "non-user" group. this makes drug users look artificially better — they appear to survive longer, but only because they had to survive long enough to get the prescription in the first place.

**the fix:** we pick a fixed time point after diagnosis (the "landmark" — e.g. 6 months). we then:

1. **exclude** all patients (users and non-users) who died before that point
2. **classify** drug exposure based only on prescriptions received before the landmark
3. **measure survival** from the landmark time forward, not from diagnosis

this ensures both groups (users and non-users) survived at least to the landmark, making the comparison fair. for als, where a meaningful fraction of patients die in the first few months, even a 6-month landmark makes a significant difference.

**interpreting results:** a hazard ratio (hr) below 1.0 means drug users survived longer from the landmark onward than matched controls. a hr above 1.0 means they survived shorter. wide confidence intervals (e.g. 0.01–60.80) indicate very small sample sizes — treat these results with caution.

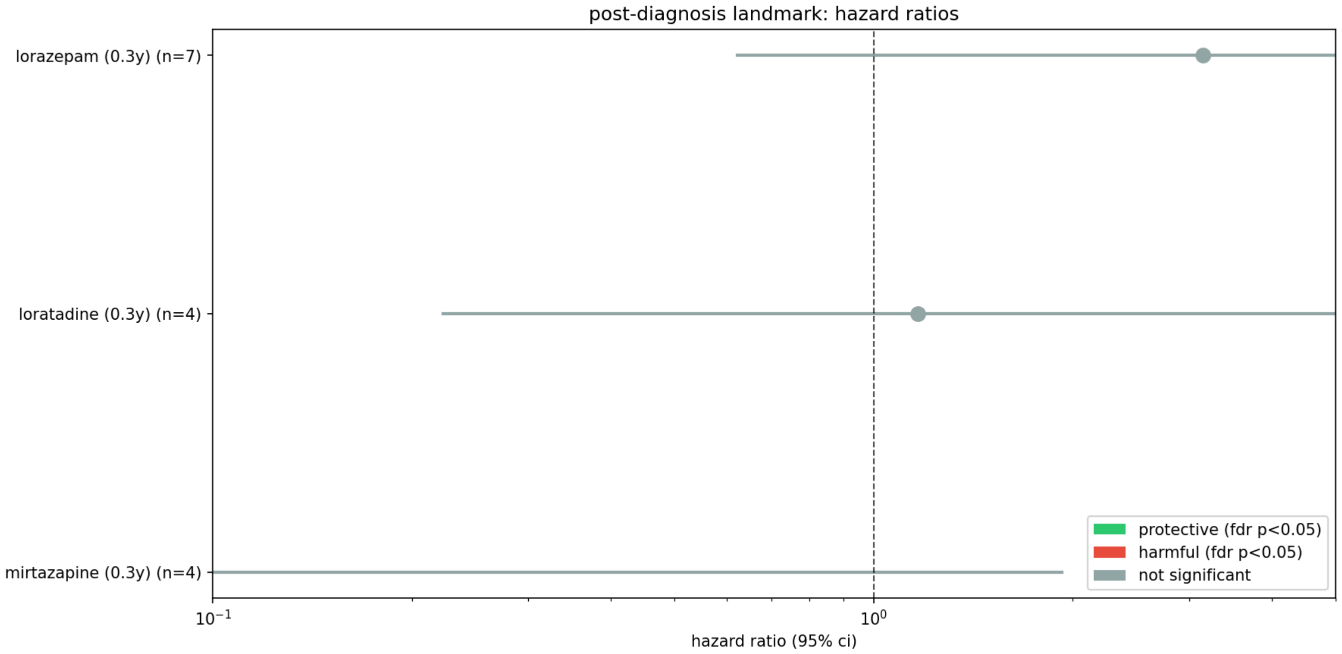


| drug | n_users | n_controls | hazard_ratio | hr_ci | logrank_pvalue |
|---|---|---|---|---|---|
| lorazepam (0.3y) | 7 | 32 | 3.148 | 0.62 – 15.92 | 0.5917 |
| mirtazapine (0.3y) | 4 | 20 | 0.039 | 0.00 – 1.93 | 0.1320 |
| loratadine (0.3y) | 4 | 20 | 1.166 | 0.22 – 6.10 | 0.0999 |

#### lorazepam

**landmark 0.3 years** — only patients who survived at least 0.3 years after diagnosis are included. survival is measured from the 0.3-year mark forward.

hr = 3.15 (0.62 – 15.92), log-rank p = 0.5917, 7 drug users vs 32 matched controls.

*drug users did not survive longer from the 0.3-year landmark (hr ≥ 1).*

*⚠ very small sample size — interpret with extreme caution. the wide confidence interval reflects substantial uncertainty.*

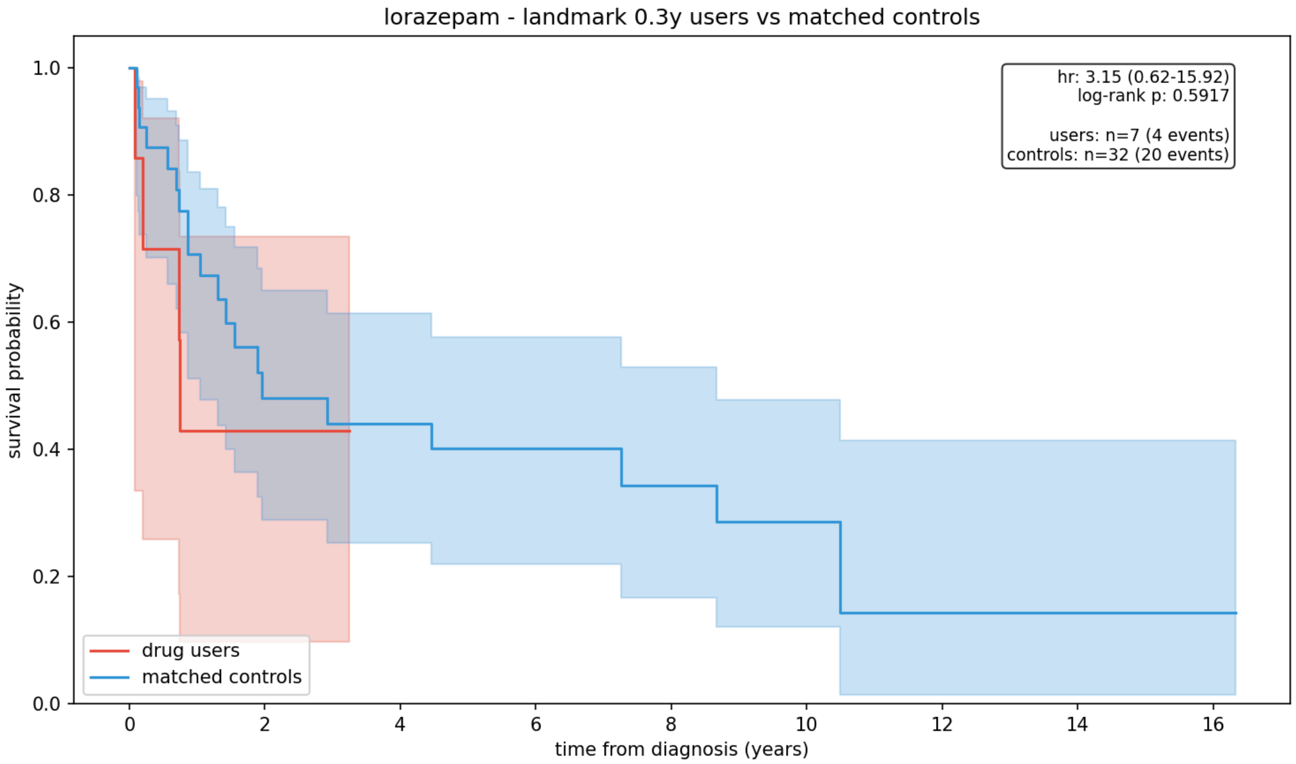


##### mirtazapine

**landmark 0.3 years** — only patients who survived at least 0.3 years after diagnosis are included. survival is measured from the 0.3-year mark forward.

hr = 0.04 (0.00 – 1.93), log-rank p = 0.1320, 4 drug users vs 20 matched controls.

*drug users survived longer from the 0.3-year landmark (hr < 1). however, this is not statistically significant (p ≥ 0.05).*

*⚠ very small sample size — interpret with extreme caution. the wide confidence interval reflects substantial uncertainty.*

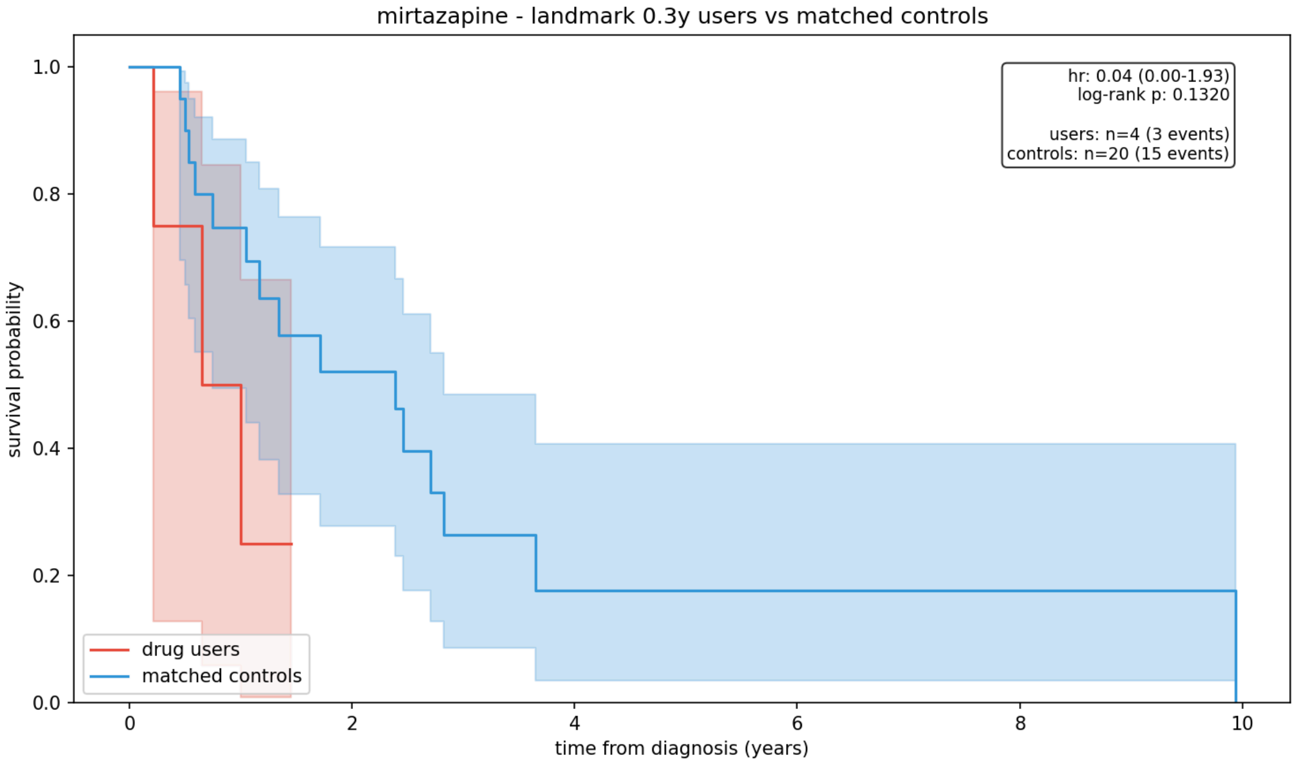

##### loratadine

**landmark 0.3 years** — only patients who survived at least 0.3 years after diagnosis are included. survival is measured from the 0.3-year mark forward.

hr = 1.17 (0.22 – 6.10), log-rank p = 0.0999, 4 drug users vs 20 matched controls.

*drug users did not survive longer from the 0.3-year landmark (hr ≥ 1).*

*⚠ very small sample size — interpret with extreme caution. the wide confidence interval reflects substantial uncertainty.*

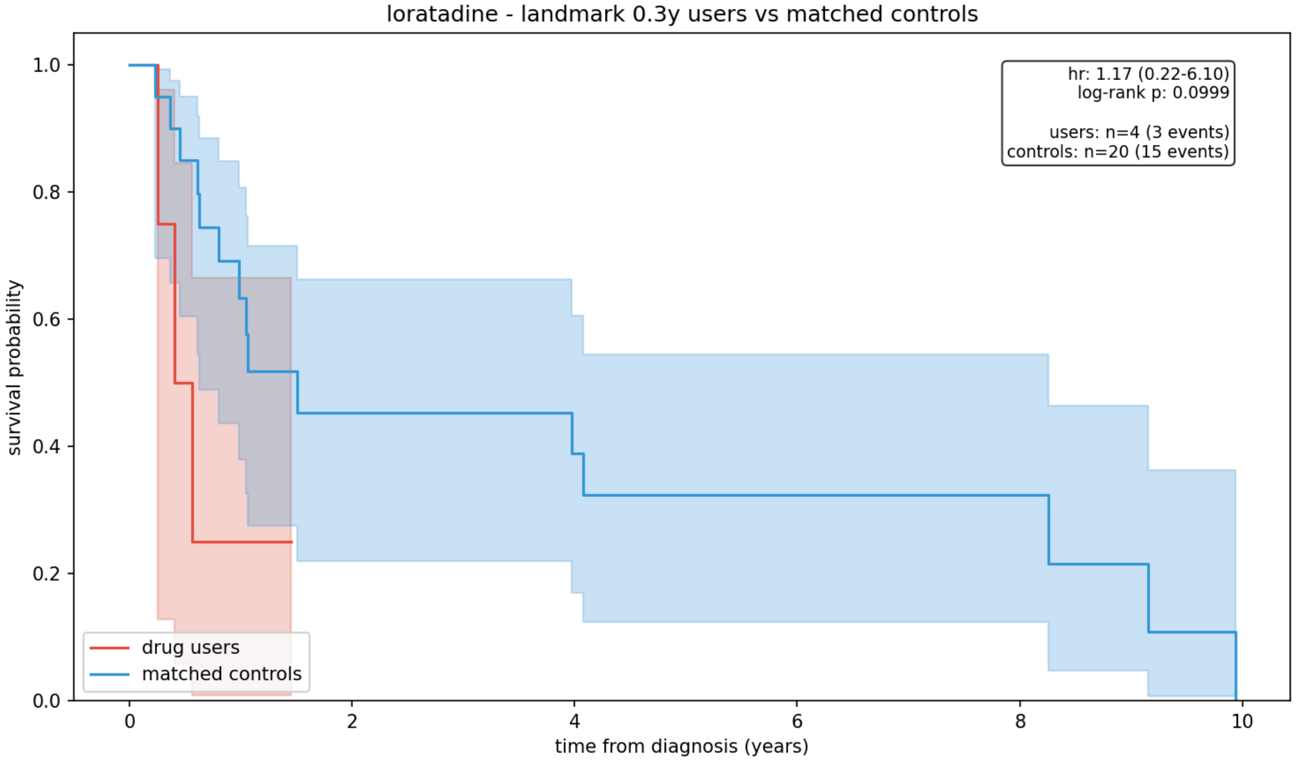


#### ATC class analysis

results grouped by ATC (Anatomical Therapeutic Chemical) classification, an international standard for organizing drugs by therapeutic use.

##### pre-diagnosis

| ATC code | n_drugs | n_users | n_controls | hazard_ratio | hr_ci | logrank_p |
|---|---|---|---|---|---|---|
| C09C | 3 | 26 | 130 | 1.024 | 0.58-1.82 | 0.3638 |
| G04C | 1 | 15 | 75 | 0.516 | 0.24-1.09 | 0.2790 |
| N05A | 2 | 4 | 20 | 5.426 | 1.03-28.47 | 0.3590 |
| N05B | 1 | 9 | 45 | 2.671 | 0.99-7.18 | 0.1080 |
| N06A | 2 | 6 | 30 | 0.247 | 0.07-0.91 | 0.6108 |
| R06A | 1 | 6 | 30 | 0.756 | 0.27-2.09 | 0.7642 |

##### post-diagnosis (landmark)

| ATC code | n_drugs | n_users | n_controls | hazard_ratio | hr_ci | logrank_p |
|---|---|---|---|---|---|---|
| N05B | 1 | 7 | 32 | 3.148 | 0.62-15.92 | 0.5917 |
| N06A | 1 | 4 | 20 | 0.039 | 0.00-1.93 | 0.1320 |

| ATC code | n_drugs | n_users | n_controls | hazard_ratio | hr_ci | logrank_p |
|---|---|---|---|---|---|---|
| R06A | 1 | 4 | 20 | 1.166 | 0.22-6.10 | 0.0999 |

#### indication-matched analysis

**what is confounding by indication?**

the biggest threat to observational drug repurposing studies is that drugs are prescribed for a *reason*. for example, if patients taking mirtazapine (an antidepressant) appear to survive longer, is that because mirtazapine is neuroprotective, or because depressed patients tend to be diagnosed earlier (lead-time bias), or because depression correlates with some other factor that affects survival?

**the fix:** indication-matched analysis restricts the comparison to patients who all have the same underlying condition. for mirtazapine, we would compare mirtazapine users vs non-users *only among patients with a depression diagnosis*. this way, both groups have the same indication — the only difference is whether they received that specific drug.

**interpreting results:** these are generally more trustworthy than the main pre-diagnosis results because they control for the reason the drug was prescribed. however, sample sizes are often much smaller (we are restricting to a subset of patients), so statistical power is reduced.

##### digoxin

insufficient data (indication patients: 9)

##### mirtazapine

insufficient data (indication patients: 0)

##### lorazepam

insufficient data (indication patients: 0)
